# DNA-PK and the TRF2 iDDR inhibit MRN-initiated resection at leading-end telomeres

**DOI:** 10.1101/2023.03.06.531339

**Authors:** Logan R. Myler, Beatrice Toia, Cara K. Vaughan, Kaori Takai, Andreea M. Matei, Peng Wu, Tanya T. Paull, Titia de Lange, Francisca Lottersberger

## Abstract

Telomeres replicated by leading-strand synthesis lack the 3’ overhang required for telomere protection. Surprisingly, resection of these blunt telomere is initiated by the telomere-specific 5’ exonuclease Apollo rather than the Mre11-Rad50-Nbs1 (MRN) complex, the nuclease that acts at DNA breaks. Without Apollo, leading-end telomeres undergo fusion, which, as demonstrated here, are mediated by alternative End Joining. Here, we show that DNA-PK and TRF2 coordinate the repression of MRN at blunt telomeres. DNA-PK represses an MRN-dependent long range resection at blunt telomeres, while the endonuclease activity of MRN/CtIP, which could cleave DNA-PK off of blunt telomere ends, is inhibited *in vitro* and *in vivo* by the iDDR of TRF2. AlphaFold-Multimer predicts a conserved association of the iDDR with Rad50 potentially interfering with CtIP binding and MRN endonuclease activation. We propose that repression of MRN-mediated resection is a conserved aspect of telomere maintenance and represents an ancient feature of DNA-PK and the iDDR.

## Introduction

TRF2 protects mammalian telomeres by forming the t-loop structure in which the 3’ singlestranded (ss) overhang invades the duplex part of the telomeres. This architectural change in the telomeric DNA has been proposed to prevent ATM kinase signaling by denying the Double Strand Break (DSB) sensor of the ATM pathway, MRN, access to the telomere end. In addition, the t-loop structure is proposed to prevent the loading of DNA-PK (comprised of the Ku70/80 heterodimer and DNA-PKcs) onto telomere ends, rendering telomeres impervious to classical Non Homologous End Joining (c-NHEJ) ^1^.

T-loop formation requires the presence of a 3’ overhang. However, after DNA replication, telomeres duplicated by leading-strand DNA synthesis are presumably blunt and require 5’ end resection to regain the 3’ overhang. At DSBs, resection is initiated by MRN/CtIP, whose endonuclease activity nicks the 5’ strand at ~15-45 nts from the break and then uses the 3’ exonuclease activity of MR to generate a short overhang ^2^. This initial resection is required for long range resection by the Exo1 exonuclease as well as DNA2, which digests 5’ ssDNA generated by the WRN or BLM RecQ helicases. Nonetheless, MRN/CtIP is not used for 3’ overhang generation at telomeres, possibly to avoid MRN-dependent ATM activation, raising the fundamental question of how MRN is kept inactive at newly replicated blunt telomeres. At budding yeast telomeres, the resection function, and possibly the checkpoint function, of the MRN ortholog, MRX, is inhibited by the interaction of the telomeric Rif2 protein with Rad50 ^3–5^. Rif2 is only found in some species of budding yeast, but, interestingly, TRF2 carries an unrelated Rad50-binding module, the iDDR (inhibitor of the DNA Damage Response) motif ^6^. At dysfunctional telomeres the iDDR minimizes the accumulation of the DNA damage factor 53BP1 ^6^, but its role at functional telomeres has not been established.

In amniotes, the 5’ exonuclease Apollo/DCLRE1B/SNM1B has evolved a YxLxP motif in its C-terminus that allows it to bind to the TRFH domain of TRF2 through the interaction with a region surrounding F120 ^7–11^. TRF2-bound Apollo is thought to initiate 5’ end resection at leading-end telomeres to allow subsequent long-range resection by Exo1 ^12^. When Apollo is deleted or prevented from binding to TRF2 (e.g., when TRF2-F120A is used to complement deletion of TRF2), leading-end telomeres do not regain their normal 3’ overhangs and are vulnerable to end joining ^13–17^. Such leading-end telomere fusions are abolished by deletion of Ku70, which initially suggested that they are mediated by c-NHEJ ^13,15^. In addition, Apollo-deficiency leads to activation of ATM signaling at a subset of telomeres, presumably the leading-end telomeres ^16,17^. Since ATM signaling at telomeres is strictly dependent on MRN ^18–21^, MRN must be associated with the unprocessed leading-end telomeres.

Therefore, leading-end telomeres deprived of Apollo are a powerful tool to investigate how MRN can activate ATM signaling but does not initiate resection at blunt telomere ends. Here we show that the answer lies in the iDDR domain of TRF2, which prevents MRN/CtlP-dependent resection in vivo and inhibits its endonuclease activity in vitro while having no effect on the MRN exonuclease activity. AlphaFold-Multimer modeling suggests that the inhibition is due to interaction of iDDR with the ATPase domain of Rad50, a mechanism analogous to the inhibition of MRX by Rif2. AlphaFold modeling also suggests that the iDDR-Rad50 interface overlaps with the binding site of CtIP, possibly explaining the inhibition of the CtIP-dependent endonuclease activity of MRN but not its exonuclease activity. We also show that resection at leading-end telomeres lacking Apollo is inhibited by DNA-PK, such that in absence of DNA-PK, compensatory resection results in protected telomeres that do not undergo fusion. This result indicates that the previously noted dependence of telomere fusions on Ku70 is not related to the role of DNA-PK in c-NHEJ but due to its ability to prevent resection when Apollo is absent. In agreement, we find that the telomere fusions in cells lacking Apollo are mediated by alt-EJ and are independent of c-NHEJ factor Ligase IV (Lig4).

## Results

### Alt-EJ mediates fusion of leading-end telomere lacking Apollo

In agreement with previous reports that the leading-end telomere fusions do not occur at blunt newly replicated leading end telomeres in cells lacking Ku70 ^13,15^, no telomere fusions were induced upon Cre-mediated deletion of Apollo from cells lacking Ku70, DNA-PKcs, or both (Fig. 1a,b and Extended Data Fig. 1a-c). Although these results could be interpreted to mean that the leading-end telomeres are joined by c-NHEJ, this is not the case since the fusions were not dependent on Lig4 (Fig. 1c,d).

**Fig. 1:**
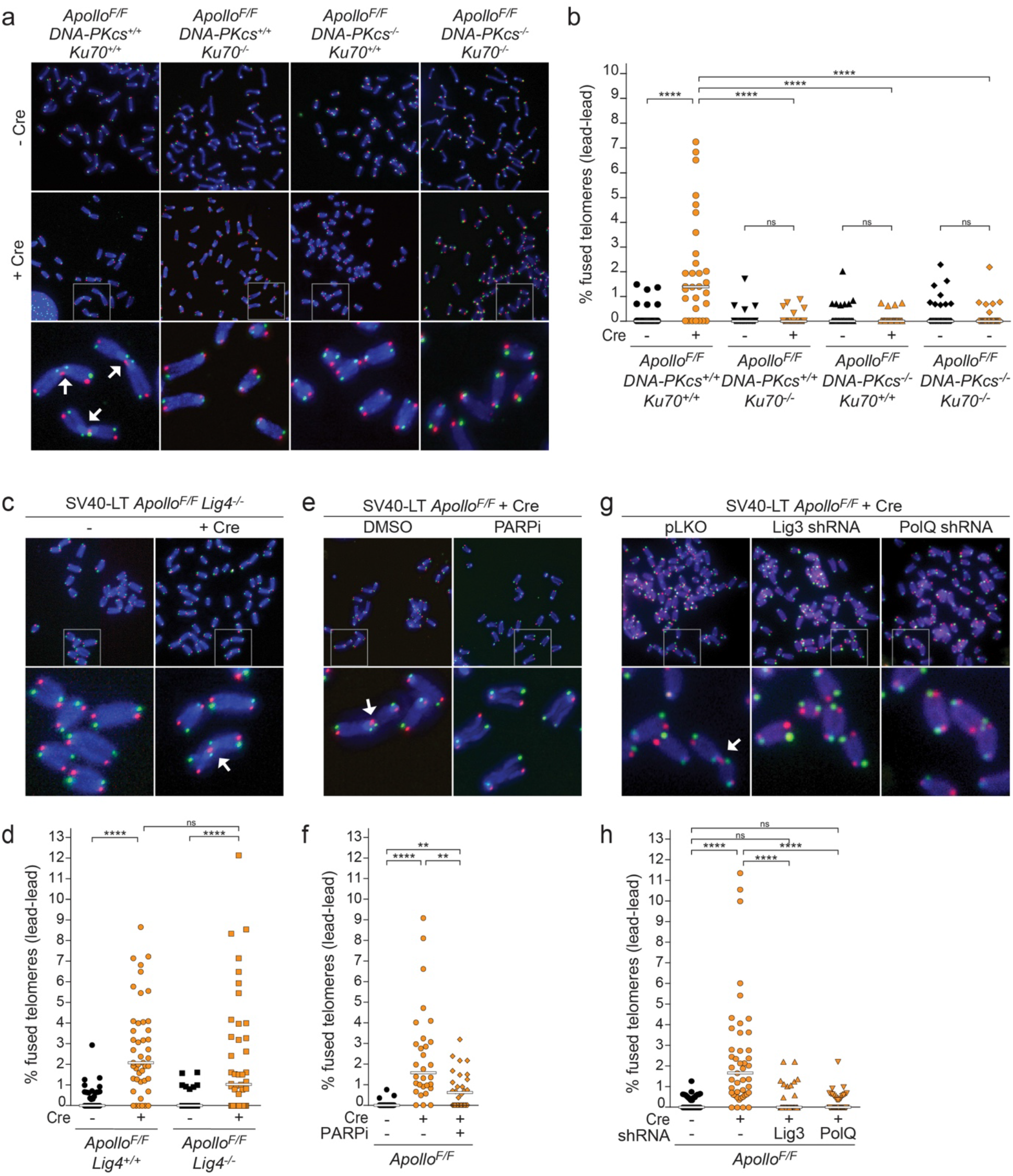
alt-EJ promotes leading-end telomere fusions due to Apollo deletion. (a) CO-FISH of metaphase spreads in SV40LT-immortalized Apollo^F/F^, Apollo^F/F^ DNA-PKcs^-/-^, Apollo^F/F^ Ku70^-/-^ or Apollo^F/F^ DNA-PKcs^-/-^ Ku70^-/-^ MEFs without any treatment or 96 h after Hit & Run Cre-mediated Apollo deletion. Leading and lagging-end telomeres were detected with Cy3-(TTAGGG)3 (red) and Alexa488-(CCCTAA)3 (green) probes, respectively. DNA was stained with DAPI (blue). Arrows indicate leading-end telomere fusions. (b) Quantification of leading-end telomere fusions as shown in (a). Each dot represents the percentage of telomeres fused in one metaphase. Bars represent the median of fused telomeres in 30 metaphases over three independent experiments. Only fusions involving two leading-end telomeres (lea/lea) are shown. Statistical analysis by Kruskal-Wallis one-way ANOVA for multiple comparisons. (c), (d) Representative metaphases and quantification of leading-end telomere fusions in Apollo^F/F^ Lig4^+/+^ and Apollo^F/F^ Lig4^-/-^ MEFs 96 hrs before and after Hit & Run Cre for three independent experiments (45 metaphases). (e), (f) Representative metaphases and quantification of leading-end telomere fusions for ApolloF/F MEFs 96 h after Hit & Run Cre and/or after 24 hrs treatment with 2 μM PARP inhibitor Olaparib (PARPi). for three independent experiments (30 metaphases). (g), (h) Representative metaphases and quantification of leading-end telomere fusions for ApolloF/F MEFs transduced with the empty vector or shRNAs agains Lig3 or PolQ and 108 h after Hit & Run Cre for three independent experiments (45 metaphases). See also Extended Data Fig. 1.

We therefore tested the role of alt-EJ in the telomere fusion events. Olaparib inhibition of PARP1 and −2, which mediate the early steps of alt-EJ ^22^, resulted in a significant reduction in the telomere fusions induced by Apollo deletion (Fig. 1e,f). Similarly, shRNAs to Ligase 3 (Lig3)^23^ or DNA polymerase theta (PolQ) caused a significant decrease in telomere fusions in cells lacking Apollo (Fig. 1g,h, Extended Data Fig. 1d,e). These data demonstrate that alt-EJ, rather than c-NHEJ, is a major mode of blunt telomere fusions and support previous reports indicating that, unlike c-NHEJ, alt-EJ can engage telomeres despite the presence of TRF2 ^24,25^.

### Loss of DNA-PK restores overhang generation at telomeres lacking Apollo

The finding that alt-EJ is responsible for the fusion of blunt telomere ends raised the question of why these fusions are not observed in absence of DNA-PK. As telomere fusions in Apollo-deficient cells are thought to be a consequence of a resection defect, we explored the possibility that loss of DNA-PK unleashes compensatory resection that restores the 3’ overhang. A role for DNA-PK as a repressor of resection would be in line with Ku70/80 blocking resection in yeast ^1–4^. Indeed, after Apollo deletion, cells lacking Ku70 or DNA-PKcs did not show the decrease in the overhang signal observed in DNA-PK-proficient cells (Fig. 2a,b). In contrast, deletion of Lig4 did not have this effect and cells lacking Lig4 showed the expected reduction in the 3’ overhang signal upon Apollo deletion, consistent with the appearance of leading-end telomere fusions (Fig. 2c,d). Conditional deletion of Ku70 from otherwise wild type MEFs did not alter the 3’ overhang signal (Figure 2e,f and Extended Data Fig. 2a,b), indicating that the effect of Ku70 on telomere resection is only apparent when Apollo is absent. Thus, DNA-PK presence appears to inhibit Apollo-independent resection at newly replicated leading-end telomeres. Without this inhibition by DNA-PK, 3’ overhang formation and telomere protection are generated independently of Apollo.

**Fig. 2:**
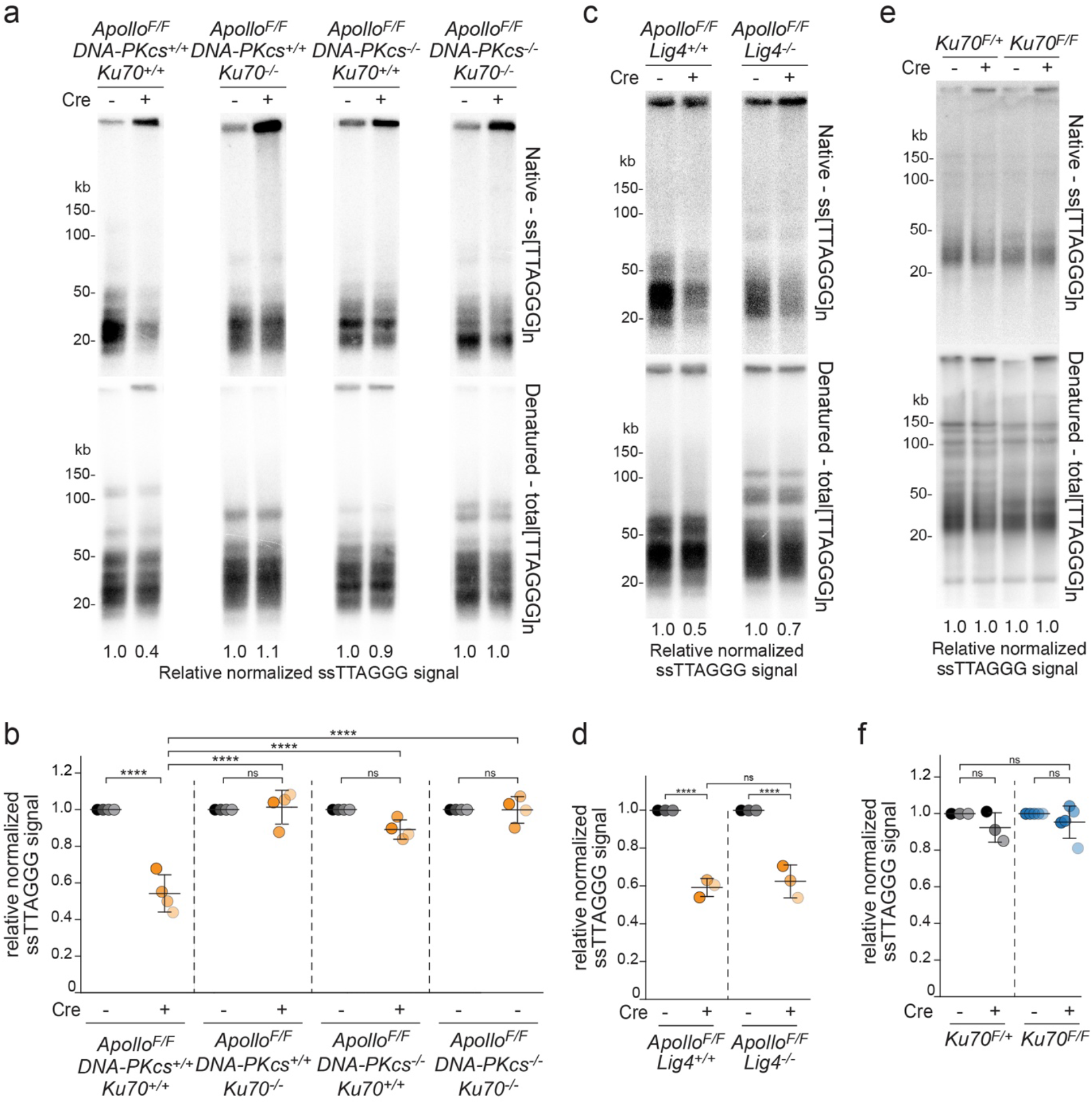
DNA-PK prevents Apollo-independent processing of 3’ telomere overhang. (a) Telomeric overhang assay on SV40LT-immortalized Apollo^F/F^, Apollo^F/F^ DNA-PKcs^-/-^, Apollo^F/F^ Ku70^-/-^ or Apollo^F/F^ DNA-PKcs^-/-^ Ku70^-/-^ MEFs, 96 h after Hit & Run Cre-mediated deletion of endogenous Apollo. Upper panel: single-stranded telomeric DNA. Lower panel: total telomeric signal. The ssTTAGGG signal was normalized to the total telomeric DNA in the same lane. The normalized no Cre value for each cell line was set to 1, and the + Cre value was given relative to it. (b) Quantification of the relative overhang signal as detected in (a) for four independent experiments (indicated by different shades), with means and SDs. Statistical analysis by two-way ANOVA. (c), (d) Telomeric overhang assay and quantification on SV40-LT-immortalized Apollo^F/F^ Lig4^+/+^ and Apollo^F/F^ Lig4^-/-^ MEFs 96 h after Cre-mediated deletion of endogenous Apollo for three independent experiments. (e), (f) Telomeric overhang assay and quantification of Ku70^F/+^ and two independent Ku70^F/F^ MEFs 96 h after Hit & Run Cre-mediated deletion of endogenous Ku70 across three independent experiments. See also Extended Data Fig. 1,2.

### NBS1 represses telomere fusions in the absence of DNA-PK and Apollo

To test how DNA-PK represses resection at telomeres, we targeted NBS1 with CRISPR/Cas9 (Fig. 3a). Bulk targeting of NBS1 did not induce telomere fusions in DNA-PK null MEFs expressing Apollo (Fig. 3b,c). However, when DNA-PKcs and Apollo were both absent, NBS1 targeting increased the frequency of leading-end telomere fusions (Fig. 3b,c), suggesting that DNA-PK represses resection by MRN. The frequency of telomere fusions induced by CRISPR/Cas9 targeting of NBS1 in DNA-PKcs null cells was lower than that observed in DNA-PK proficient Apollo KO cells (Fig. 3b,c). Although it is possible that the CRISPR/Cas9 targeting was not sufficient to abort MRN-dependent resection in all cells, there are other explanations that are discussed below.

**Fig. 3:**
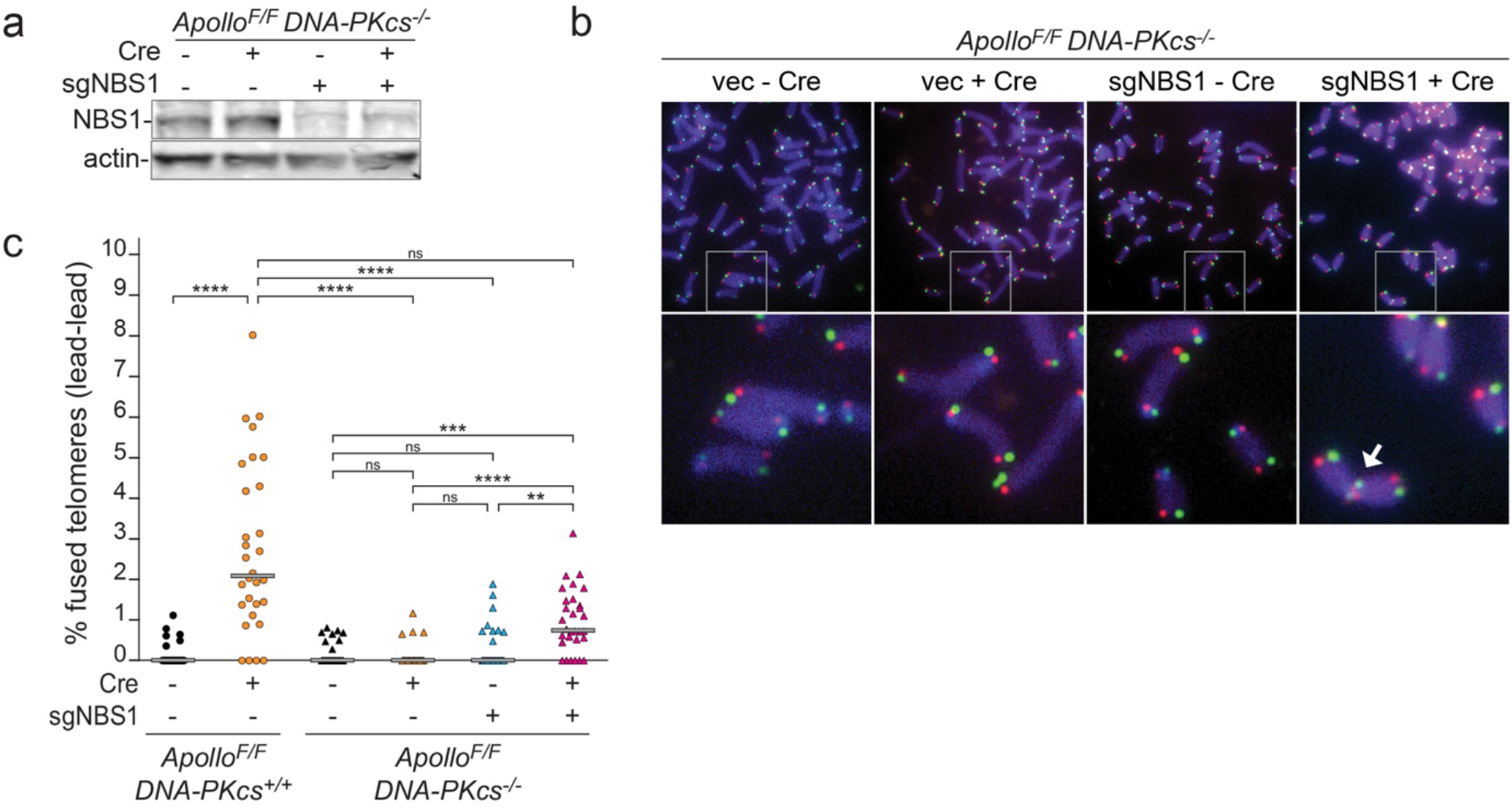
NBS1 protects from leading-end telomeres fusions in the absence of Apollo and DNA-PKcs. (a) Immunoblots for NBS1 in SV40LT-immortalized Apollo^F/F^ DNA-PKcs^-/-^ MEFs, after transduction with Cas9 expressing vector with or without sgRNA against NBS1 and/or Hit & Run Cre. Actin is shown as loading control. (b) and (c) CO-FISH metaphase analysis and quantification of leading-end telomere fusions in Apollo^F/F^ and Apollo^F/F^DNA-PKcs^-/-^ MEFs 96 h after Cre-mediated deletion of Apollo and/or 48 h after deletion of NBS1 by CRISPR/Cas9 and specific sgRNA, as shown in (a). Graph represents 30 metaphases over three independent experiments. Statistic by by Kruskal-Wallis one-way ANOVA for multiple comparisons.

### The iDDR of TRF2 inhibits MRN/CtIP at newly repliated leading-end telomeres

The observation of DNA-PK preventing resection at blunt telomeres is surprising, given that DNA-PK promotes MRN/CtIP endonuclease activity at DSBs^29^, and it suggests that MRN/CtIP has different access to telomeres than DSBs. As the iDDR of TRF2 interacts with MRN ^6^, we asked whether the iDDR affects MRN-initiated resection at telomeres lacking Apollo. We complemented conditional TRF2^F/F^ cells with wild type TRF2 (WT), the TRF2ΔiDDR allele (ΔiDDR), the TRF2-F120A allele (F120A) that does not bind Apollo ^7^, or a version of TRF2 containing both mutations (F120A-ΔiDDR) (Fig. 4a and Extended Data Fig. 3a,b). Importantly, removal of the iDDR completely mitigated the effect of Apollo loss, restoring the 3’ overhang signal and abolishing leading-end telomere fusions (Fig. 4b-e). In addition, the deletion of the iDDR by itself caused a small reduction in proliferation and an increase of g-H2AX foci at telomeres as well as a trend toward greater telomeric overhang signals for reasons that remain to be determined (Fig. 4b,c and Extended Data Fig. 3c-e).

**Fig. 4:**
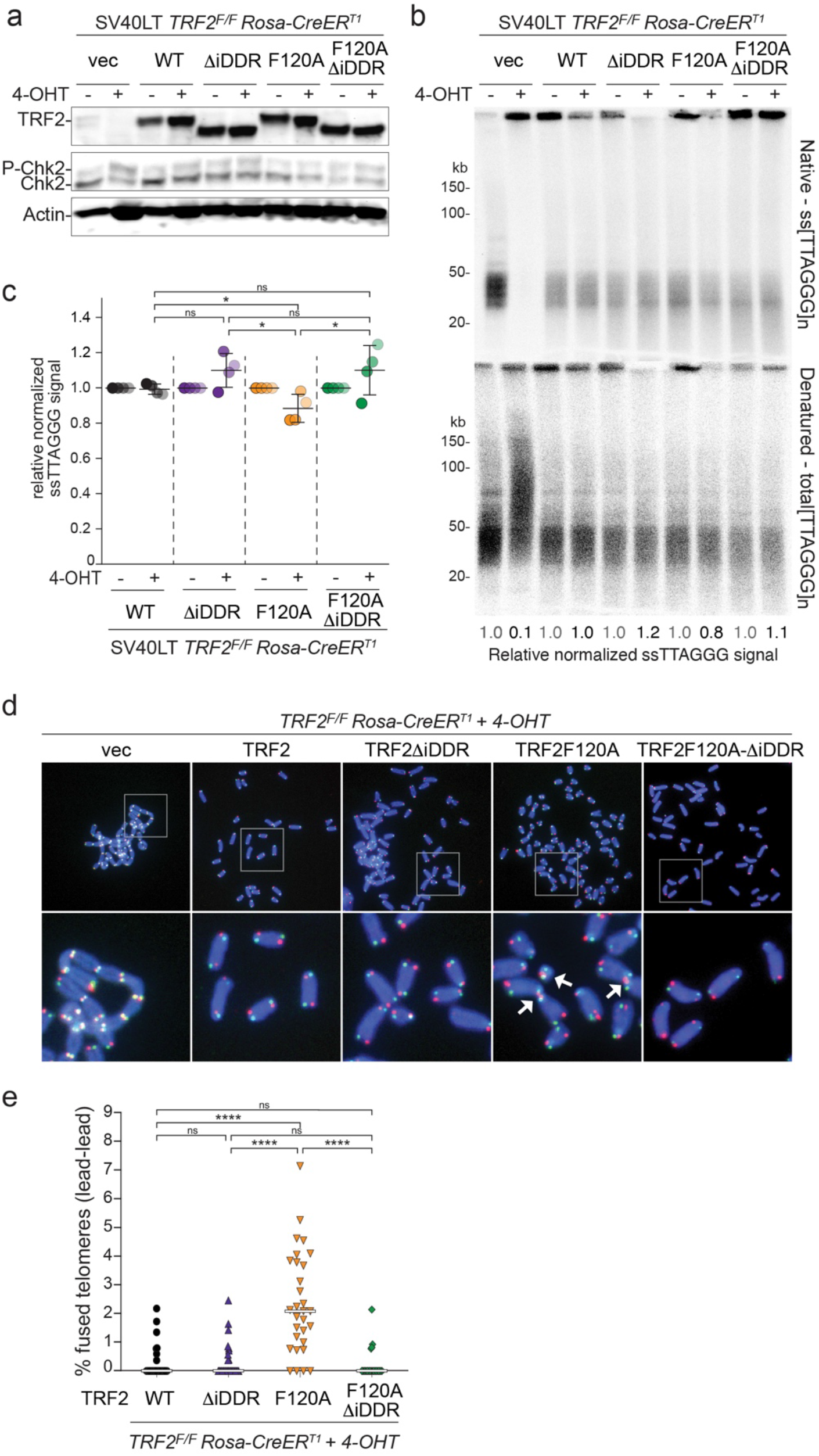
TRF2 prevents Apollo-independent processing of 3’ telomere overhang through the iDDR region. (a) Immunoblot of endogenous and exogenous TRF2 and Chk2 phosphorylation in SV40LT-immortalized TRF2^F/F^ RsCre-ERT1 MEFs expressing the empty vector (EV), or the MYC-TRF2 (WT), MYC-TRF2-ΔiDDR (ΔiDDR), MYC-TRF2-F120A (F120A) or MYC-TRF2-F120AΔiDDR (F120AΔiDDR) alleles at 96 h after 4-OH tamoxifen (4-OHT)-mediated deletion of endogenous TRF2. Actin is shown as loading control. (b) and (c) Telomeric overhang assay and quantification from three independent experiments of cells treated as described in (a). For each MYC-TRF2 allele the normalized no Cre value was set to 1, and the + Cre value was given relative to it, with means and SDs. Statistical analysis by unpaired t-test. (d) and (e) CO-FISH metaphase analysis and quantification of leading-end telomere fusions on TRF2^F/F^ RsCre-ERT1 MEFs expressing the empty vector (EV) or the MYC-TRF2 alleles described in (A) 96 h after treatment with 4-OHT. Graph represents 30 metaphases over three independent experiments. Statistic by by Kruskal-Wallis one-way ANOVA for multiple comparisons. See also Extended Data Fig. 3.

To establish whether the iDDR acts by controlling MRN, we expressed the same alleles of TRF2 in cells lacking NBS1 and examined the telomeric overhang signals and telomere fusions after TRF2 deletion. Consistent with the results presented above, in the NBS1-proficient cells, absence of the iDDR mitigated the 3’ overhang defect observed in absence of Apollo. In contrast, the deletion of the iDDR from TRF2-F120A did not improve the processing of leading-end telomeres when NBS1 was absent (Fig. 5a,b). Similarly, the leading-end fusions at telomeres lacking Apollo were diminished when the iDDR was removed from TRF2 in NBS1-proficient cells. In contrast, in NBS1-deficient cells, removal of the iDDR from TRF2 did not affect the leading-end telomere fusions due to lack of Apollo recruitment (Fig. 5c,d). These results indicate that the iDDR of TRF2 acts through MRN.

**Fig. 5:**
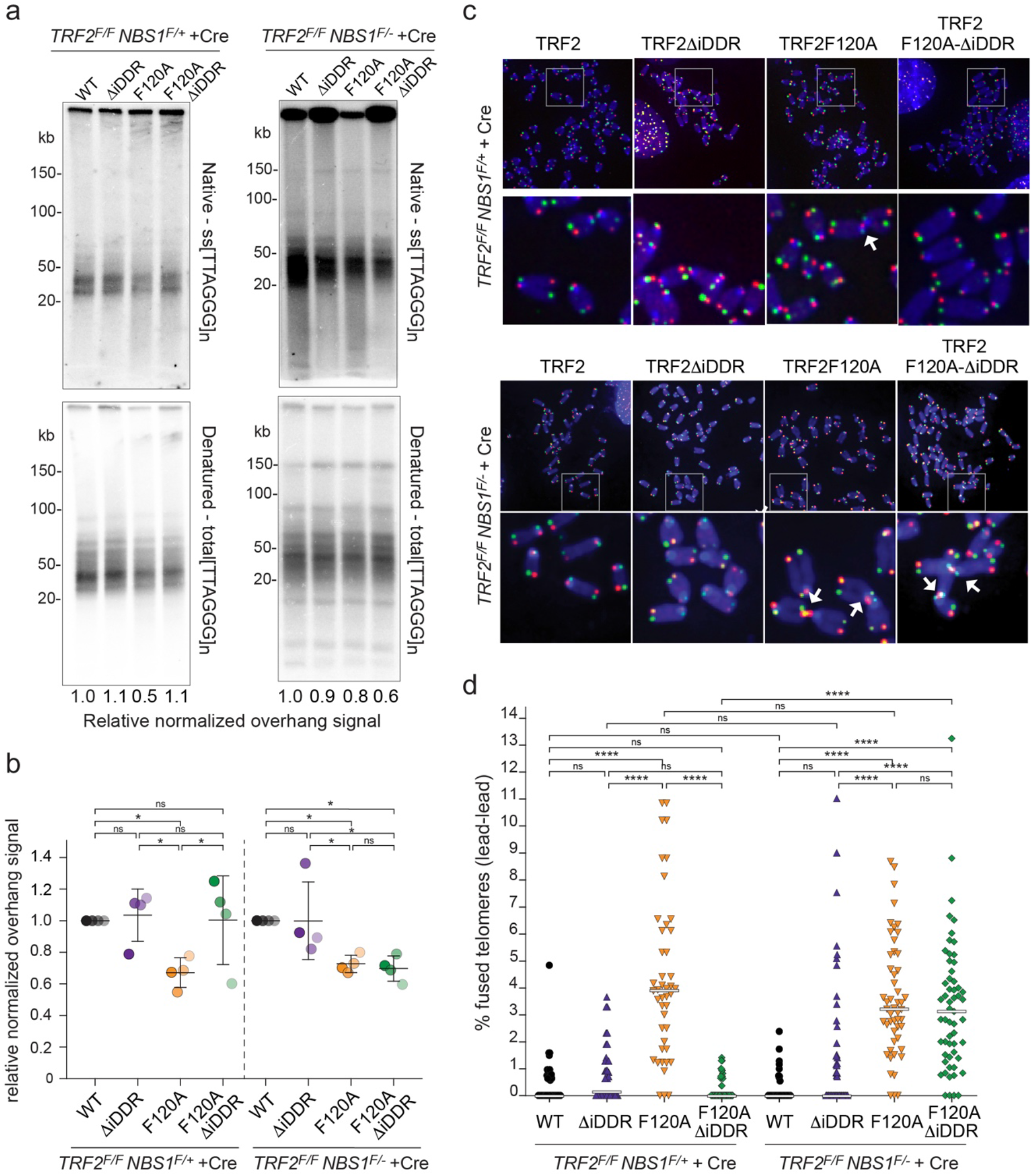
NBS1 is required for the Apollo-independent processing of 3’ telomere overhang inhibited by the TRF2 iDDR. (a) and (b) Telomeric overhang assay and quantification from three independent experiments of TRF2^F/F^NBS1^F/+^ and TRF2^F/F^NBS1^F/-^ MEFs expressing the indicated MYC-TRF2 alleles 120 hr after Cre-mediated deletion of TRF2 or TRF2 and NBS1. Each cell line was normalized to the MYC-TRF2-WT allele with means and SDs. Statistical analysis by unpaired t-test. (c) and (d) CO-FISH metaphase analysis and quantification of leading-end telomere fusions in TRF2^F/F^NBS1^F/-^ and TRF2^F/F^NBS1^F/-^ MEFs expressing the indicated MYC-TRF2 alleles 120 h after Cre-mediated deletion of TRF2 or TRF2 and NBS1. Graph represents 30 metaphases over three independent experiments with medians. Statistic by by Kruskal-Wallis one-way ANOVA for multiple comparisons. See also Extended Data Fig. 4.

### No role for 53BP1

Since 53BP1 blocks the formation of excessively long 3’ overhangs at dysfunctional telomeres ^23,30–32^, we tested whether 53BP1 also affected the formation of 3’ overhangs at telomeres lacking Apollo. However, in 53BP1-deficient cells, the complementation of TRF2 deletion with the F120A allele again caused a reduction of the 3’ overhang signal, and, as for 53BP1 proficient cells, no reduction was observed in cells complemented with the F120A-ΔiDDR allele (Extended Data Fig. 4a). Furthermore, 53BP1 status had no effect on the reduction in the 3’ overhang after Apollo deletion (Extended Data Fig. 4b,c), indicating that the iDDR inhibits MRN/CtIP independent of 53BP1.

### In vitro inhibition of MRN/CtIP endonuclease by the iDDR

Given that the iDDR inhibits MRN in a 53BP1-independent manner, we asked whether it directly affects MRN activity. MRN is an endonuclease that is activated by phosphorylated CtIP to nick the 5’ strand at protein-blocked DNA ends ^33–35^. The endonuclease activity of MRN-CtIP can be measured under physiological conditions on a DNA substrate that is bound to DNA-PK. In this setting, MRN generates an endonucleolytic product of approximately 45 nt ^29^. Indeed, purified TRF2 inhibited the formation of this product in a dose-dependent manner (Fig. 6a-c). By contrast, TRF2ΔiDDR motif did not have this effect, indicating that the iDDR is required for the inhibition MRN endonuclease activity. MRN also exhibits a 3’-5’ exonuclease activity that is independent of CtIP ^36,37^. Contrary to its role in inhibiting the endonuclease activity of MRN, the addition of TRF2 or TRF2ΔiDDR did not inhibit the exonuclease activity of MRN (Fig. 6d-f). These results show that the iDDR has a specific effect on the endonuclease activity of MRN/CtIP but does not interfere with the DNA end binding or exonuclease activity of MRN. The effect of the iDDR is similar to that of budding yeast Rif2, which inhibits MRX endo-but not exonuclease activity ^3,4^.

**Fig. 6:**
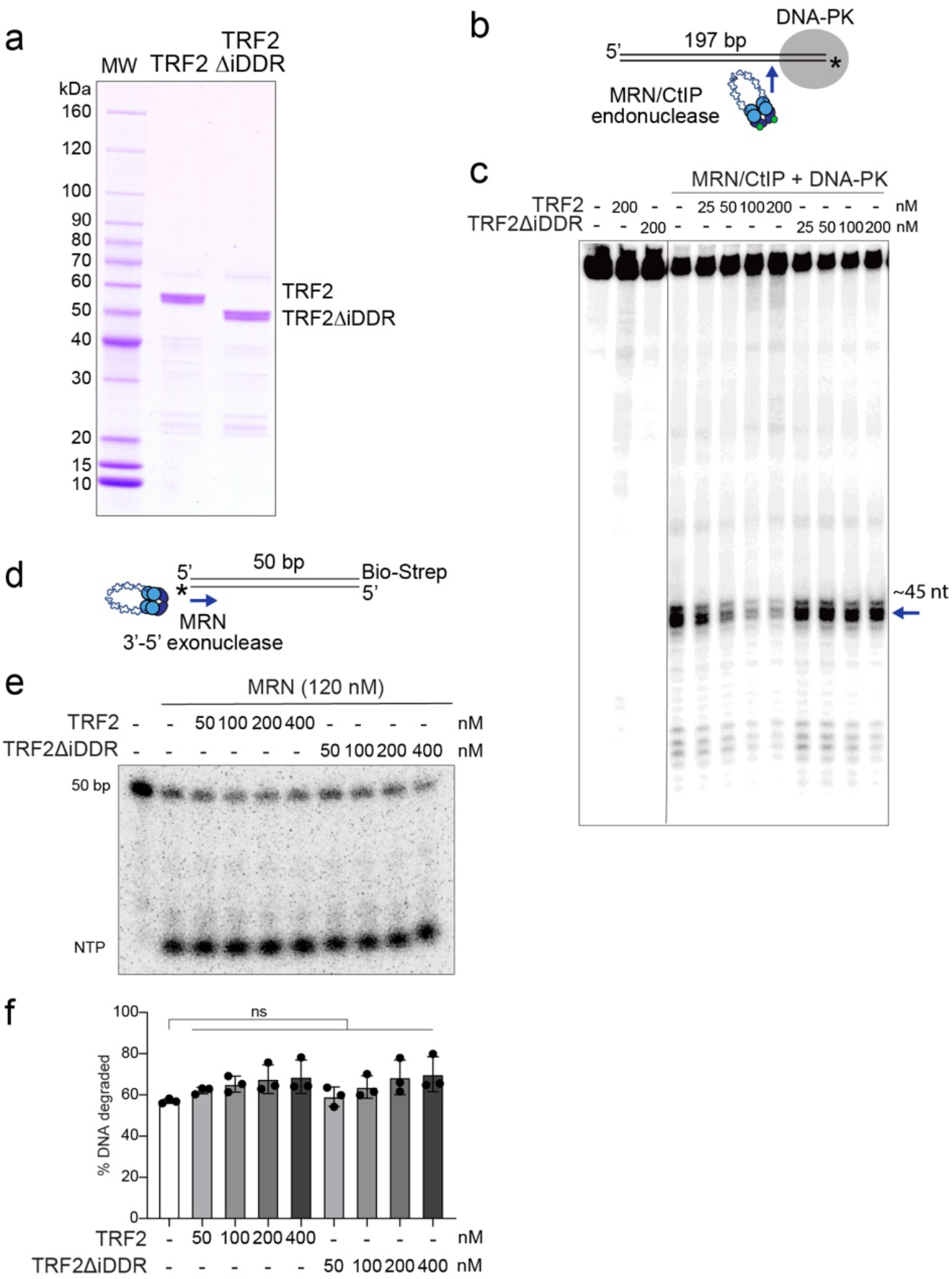
The iDDR of TRF2 inhibits MRN endonuclease activity in vitro. (a) Coomassie-stained SDS-PAGE gel of purified TRF2 proteins. (b) Schematic of the MRN/CtIP endonuclease assay with DNA-PK. (c) MRN endonuclease assay in the presence of TRF2-WT or TRF2-ΔiDDR. MRN (50 nM) was incubated with CtIP (80 nM), DNA-PKcs (10 nM), Ku (10 nM) and varying concentrations of TRF2 (25, 50, 100, or 200 nM) in the presence of a 5’ 32P-labeled DNA substrate. The gel is a representative example of two independent replicates. Blue arrow indicates primary endonucleolytic cleavage product (~45 nt away from the end). (d) Schematic of the MRN exonuclease assay. (e) and (f) MRN endonuclease assay gel and quantification of three independent replicates in the presence of TRF2-WT or TRF2-ΔiDDR. Statistics by unpaired t-test assuming a gaussian distribution.

### Alphafold-modeling predicts a conserved interaction between the iDDR and Rad50

It was previously shown that the iDDR of TRF2 pulls down Rad50 in co-immunoprecipitation experiments ^6^. However, these experiments did not determine which subunit of MRN interacts with TRF2. We queried the potential interactions of the human iDDR with MRN subunits using AlphaFold-Multimer ^38^. AlphaFold-Multimer predicted an interaction between the iDDR and the globular ATPase domain of Rad50 (Fig. 7a-c). The iDDR was predicted with high confidence (pLDDT value) in the top five ranked models (Fig. 7b). No interactions were predicted with Mre11, Nbs1, or CtIP, and the iDDR was predicted with low confidence (pLDDT value) in these models, suggesting that Rad50 binding orders the domain (Extended Data Fig. 5a). The predicted interaction of the iDDR with Rad50 is highly conserved in metazoans, including both vertebrate TRF2 and invertebrate TRF proteins (Extended Data Fig. 5b,c).

**Fig. 7:**
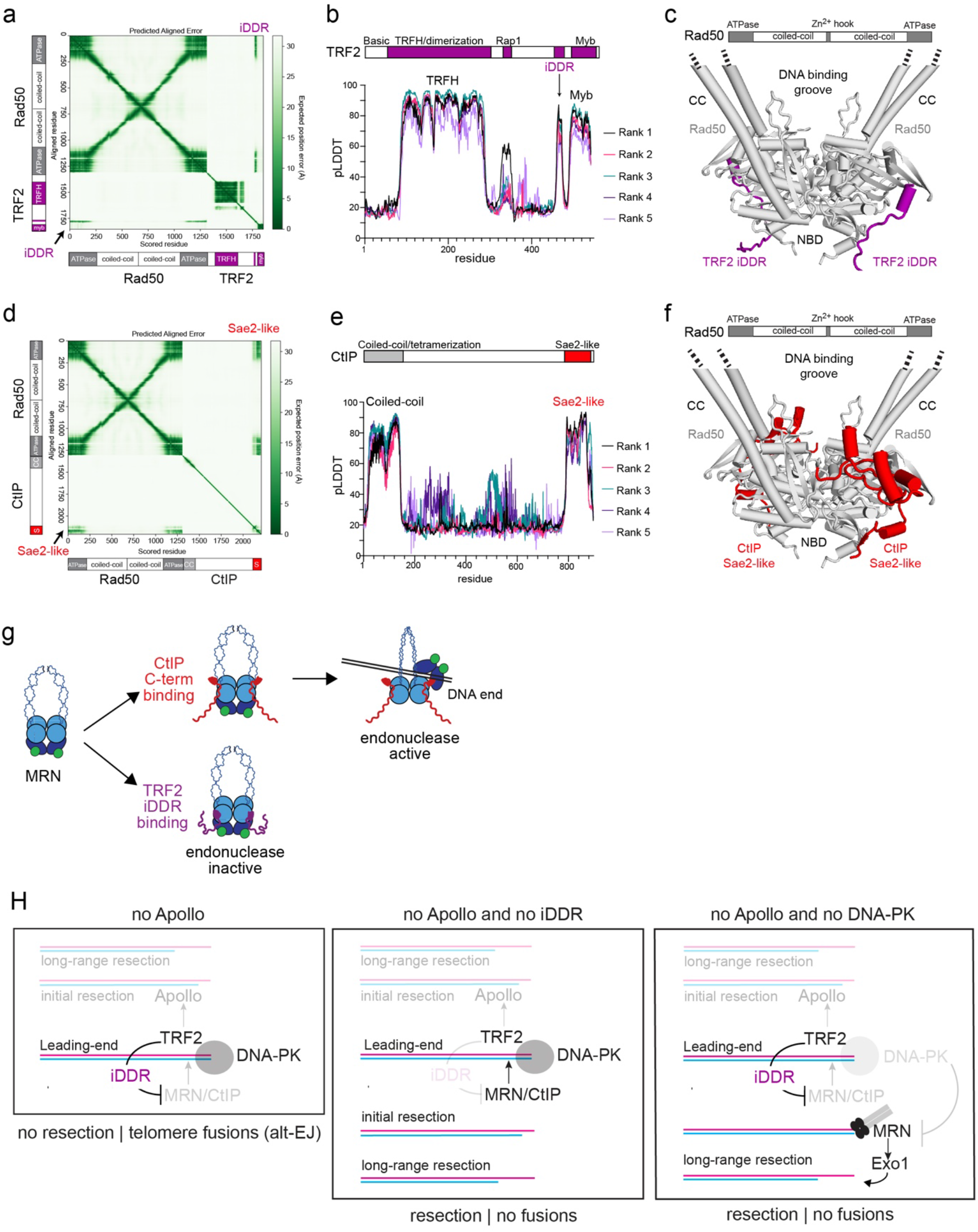
The iDDR of TRF2 is predicted to compete with CtIP for binding to Rad50. (a) Predicted Aligned Error (PAE) plot from the AlphaFold-Multimer modeling of human Rad50-TRF2. Representative of five ranked models generated with default parameters. (b) Predicted Local Distance Difference Test (pLDDT) plot showing per-residue confidence score across human TRF2 from the ranked Rad50-TRF2 models. (c) Predicted structure of a human Rad50 dimer with a dimer of human TRF2. The Rad50 coiled-coils were truncated for the dimer model. Only the iDDR of TRF2 is shown. (d) PAE plot from the AlphaFold-Multimer modeling of human Rad50-CtIP. Representative of five ranked models generated with default parameters. (e) pLDDT plot showing per-residue confidence score across human CtIP from the ranked Rad50-CtIP models. (f) Predicted structure of a human Rad50 dimer with the Sae2-like domain of human CtIP. The Rad50-CtIP monomer model was superimposed on the predicted structure of a Rad50 dimer from (c). (g) Model for inhibition of MRN endonuclease activity by the TRF2-iDDR. TRF2 competes with CtIP binding to Rad50, which stimulates the endonuclease-active state of MRN. (h) Model for leading-end telomere processing and protection mediated by TRF2 and DNA-PK. TRF2 promotes the 5’-resection of the leading-end telomere by recruiting Apollo. In the absence of Apollo, either due to Apollo deletion or the lack of recruitment due to the TRF2-F120A mutation, the newly replicated leading-end telomere ends cannot be resected and undergo fusion mediated by alt-NHEJ. In the absence of Apollo and the iDDR domain of TRF2, MRN/CtIP initiate resection at DNA-PK-bound leading end telomeres, leading to telomere protection and the absence of fusions. When both DNA-PK and Apollo are absent, MRN can promote the resection of the free DNA ends even in the presence of the TRF2 iDDR domain. See also Extended Data Fig. 5-7.

The telomeric binding proteins Rif2 and Taz1 of budding and fission yeast, respectively (Extended Data Fig. 6a), have recently been proposed to bind to Rad50 via their MRN(X)-inhibitory (MIN) domains (also called the BAT domain in Rif2) ^3,5^. AlphaFold-Multimer predicted that Rif2 and Taz1 bind their cognate Rad50 proteins using the same interface in the ATPase domain used by the iDDR (Extended Data Fig. 6b, c). Despite this analogous binding site, the MIN domains of Rif2 and Taz1do not show sequence homology and are predicted to be structurally different from the iDDR domain. In fact, the iDDR consists of a cluster of basic residues followed by acidic residues, which create a characteristic drop in local isoelectric point (pI) (Extended Data Fig. 6d, e; ^10^). On the contrary, the MIN domains of Rif2 and Taz1 have a switched order of basic and acidic residues. This inversion is reflected in their predicted orientation on the surface of Rad50 with the N-terminal residues of the MIN domain pointing toward the coiled-coils, whereas in the iDDR the C-terminus is pointing towards the coiled-coils (Extended Data Fig. 6d). This inverted orientation suggest that these motifs may have evolved independently and that their similar function is due to convergent evolution (Extended Data Fig. 6f), and, therefore, that an MRN-inhibitory module is a highly selected and ancient feature of telomeres.

### The iDDR is predicted to compete with CtIP for binding to Rad50

Data from budding yeast suggest that Rif2 inhibits the MRX endonuclease activity by outcompeting Sae2 at its Rad50 binding site ^4,33^. Therefore, we used AlphaFold-Multimer to query whether the iDDR might similarly affect the interactions between CtIP and the MRN complex. The “Sae2-like” C-terminus of CtIP was predicted with high confidence to interact with the same surface of Rad50 as the iDDR of TRF2 (Fig. 7d-f). Interestingly, this is the region of Rad50 where the Rad50S separation of function mutations cluster, which are deficient in CtIP-dependent endo-but not in the CtIP-independent exonuclease activity ^39–41^. This region of CtIP is highly-conserved across metazoans, consistent with its potential role in regulating the also highly-conserved MRN complex (Extended Data Fig. 7a). Additionally, Sae2 and Ctp1 (the fission yeast homolog of CtIP) were predicted to bind to their respective Rad50 proteins in a similar manner (Extended Data Fig. 7b-d). Collectively, the data suggest that the iDDR of TRF2 can prevent CtIP from binding to Rad50, blocking the activation of MRN endonuclease activity (Fig. 7g).

## Discussion

These results reveal the ability of the TRF2 iDDR and DNA-PK to independently inhibit all MRN-mediated resection at blunt-ended telomeres, although they resemble DSBs (Fig. 7h). When telomeres are replicated in the absence of Apollo, the leading-strand DNA synthesis products lack the protective 3’ overhang and become vulnerable to alt-EJ. These fusion events depend on the ability of DNA-PK to prevent long-range resection. However, it has been puzzling why MRN/CtIP does not promote DNA end processing by removing DNA-PK from these telomeres, as it does at DSBs. Here we show that the iDDR of TRF2 blocks MRN/CtIP endonuclease activity in vitro and prevents MRN from acting at blunt leading-end telomeres in vivo. Our structure prediction data suggest that the iDDR acts by preventing the formation of the MRN/CtIP complex. Because MRN/CtIP is not active at telomeres bearing the iDDR, DNA-PK can persist at these ends, leading to complete absence of resection. When the iDDR is removed, MRN/CtIP is active and can cleave DNA-PK off the telomere ends. Interestingly, when DNA-PK is absent, MRN can still engage the ends, leading to long-range resection, although the endonucleolytic activity of MRN/CtIP should be held in check by the iDDR. In both scenarios, resection is activated. Paradoxically, this leads to the formation of a protective 3’ overhang independently of Apollo. Importantly, we show that the inhibition of MRN endonuclease activity at eukaryotic telomeres is conserved, in part due to convergent evolution.

The iDDR is an ancient feature of the TRF subunit of shelterin. It was already present when metazoans emerged, prior to the gene duplication that created TRF1 and TRF2 and long before the Apollo-TRF2 binding evolved ^10^. In contrast, the genes involved in the ability of mammalian iDDR to minimize the accumulation of 53BP1 at dysfunctional telomeres, including BRCC3, RNF8, RNF168, and 53BP1 itself, are not conserved in all metazoans ^42^. This argues that the iDDR evolved the ability to affect 53BP1 later in evolution as a secondary feature. The argument that inhibiting MRN is the original function of the iDDR is strengthened by the finding that the MRN/X inhibitory modules (MIN/BAT) are found at telomeres in fungi. Remarkably, these fungal MIN modules interact with the same part of Rad50 as the iDDR. Because the MIN/BAT have no sequence similarity to the iDDR, are found in proteins that are not orthologous to TRF2 (e.g. Rif2), and bind in an inverse orientation compared to the iDDR, we infer that the inhibition of MRN at telomeres represents an example of convergent evolution. Such convergent evolution speaks to the strong selective pressure to preserve the inhibition of MRN at telomeres, despite major changes in the telomere-associated proteins. Why the endonuclease activity of MRN needs to be inhibited at telomeres remains to be determined.

DNA-PK blocks the access of the long-range nucleases Exo1 and DNA2/BLM to DNA ends in vitro and suppress DNA end resection in vivo ^29,43,44^. However, counter-intuitively, the presence of DNA-PK also stimulates the MRN/CtIP endonuclease activity in vitro ^29^. These results are consistent with the essential role of MRN/CtIP in initiating resection at DNA-PK blocked ends such as single-ended DSBs resulting from replication fork collapse ^45^. Our data demonstrate that the iDDR inhibits MRN/CtIP from acting as an endonuclease, effectively blocking all MRN/CtIP-mediated resection at the DNA-PK bound telomeres. Importantly, DNA-PK allows Apollo processing. However, in the absence of DNA-PK, telomeres can be processed independently of Apollo. This resection seems to be dependent on Nbs1, at least partially. Therefore, we propose that MRN can promote the resection of blunt-ended telomeres when DNA-PK is absent, possibly due to its ability to load other resection factors at the ends. We further propose that this loading activity of MRN does not require CtIP, explaining why it is not repressed by the iDDR. In agreement, MRN promotes loading of Exo1 and the other long-range resection factors at DNA ends in vitro and enhances the processivity of Exo1 resection in the presence of RPA ^46–48^. Whether other resection regulators, in addition to MRN, gain access to the telomere ends that lack DNA-PK remains to be determined.

The data show that the leading-end telomeres in cells lacking Apollo become joined by PolQ-dependent alt-EJ (also called Theta Mediated End Joining, TMEJ). This result was unexpected because leading-end telomeres lacking Apollo are presumed to be blunt whereas alt-EJ joins DSBs with 3’ overhangs. In alt-EJ, PolQ uses microhomologies to anneal 3’ overhangs and then executes templated DNA synthesis to creates the substrate for Lig3-mediated ligation. Since PolQ is auto-inhibited at DNA ends with short or no 3’ overhangs ^52^, it remains to be determined how it acts on the blunt leading-end telomeres. Conversely, it is prudent to ask why alt-EJ does not act on telomeres that have a 3’ overhang which can anneal to form 2 bp every six nucleotides. Most likely, this is due to the presence of the POT1 (POT1a and POT1b in mice) on the ssDNA since alt-EJ has been shown to act at telomeres that lack POT1 ^25,53^. Finally, it is curious that the blunt leading-end telomeres formed in absence of Apollo are not processed by c-NHEJ, despite their interaction with DNA-PK. Perhaps TRF2 inhibits c-NHEJ in S/G2 through Rap1 ^54^ or by binding to Ku70/80 ^55^. Such a S/G2-specific mechanisms of c-NHEJ repression may also account for the lack of c-NHEJ repair of telomere-internal DSBs created in S/G2 ^24^.

## Notes

### Competing Interest Statement

TdL is a member of the SAB of Calico LLC, San Francisco, CA, USA. The other authors declare no competing interest.

