## Supplemental information and figures for "DNA-PK and the TRF2 iDDR inhibit MRN-initiated resection at leading-end telomeres"

**Extended Data**

**Methods**

**Cell lines and cell treatments**

SV40-LT Apollo^F/F^, TRF2^F/F^ Rosa26 Cre-ER^T1^, TRF2^F/F^ 53BP1^-/-^ Rosa26 Cre-ER^T1^, TRF2^F/F^ NBS1^F/+^, and TRF2^F/F^ NBS1^F/-^ MEFs have been previously described ^1–3^. Apollo^F/F^ DNAPK^-/-^, Apollo^F/F^ Ku70^-/-^, Apollo^F/F^ DNAPK^-/-^ Ku70^-/-^, and Apollo^F/F^ Lig4^-/-^ were obtained by intercrosses between the respective single mutant mice ^4–6^. To generate Ku70^F/F^ mouse, the Ku70 gene (Xrcc6, chromosome 15) was modified by gene targeting. Male ES cells Xrcc6^tm1a(KOMP)Mbp^ were obtained from Knockout Mouse Programme (KOMP) and used to derive heterozygous mice. The LacZ/Neo insert was removed by crossing to flippase mouse, resulting in the floxed allele (Ku70^F^). Standard crosses of Ku70^F/+^ mice were used to derive Ku70^F/F^ MEFs. All MEFs were isolated from E12.5 or E13.5 embryos, immortalized at passage 2 with pBabeSV40LargeT (a gift from G. Hannon), and cultured in Dulbecco’s Modified Eagle Medium (DMEM) (Cellgro) supplemented with 15% fetal bovine serum (FBS) (Gibco), non-essential amino acids (Gibco), L-glutamine (Gibco), penicillin/streptomycin (Gibco), 50 μM β-mercaptoethanol (Sigma). Genotyping was performed by Transnetyx Inc. using real-time PCR. 293T and Phoenix eco cells (ATCC, Rockville, MD) were cultured in Dulbecco’s Modified Eagle Medium (DMEM) (Corning) supplemented with 10% HyClone Calf Serum (Cytiva), non-essential amino acids (Gibco), L-glutamine (Gibco), penicillin/streptomycin (Gibco). PARP inhibitor (PARPi) (Olaparib, AZD2281/ KU-59436, BioVision) was dissolved in DMSO and added at a final concentration of 2 μM for 24 h. Rosa26 Cre-ER^T1^ was induced by incubation with 1 μM 4-OHT for 24 hours.

**Viral gene delivery**

For retro or lentiviral transduction, a total 20 μg of plasmid DNA was transfected into Phoenix eco or 293T cells, respectively, using CaPO_4_ precipitation. The viral supernatant was filtered through a 0.45-μm filter, supplemented with 4 μg/ml polybrene and used for the transduction of target cells. Cre was induced with three infections/day (6-12 h intervals) over two days with pMMP Hit & Run Cre retrovirus produced in Phoenix eco cells or by adding 1 μM 4-OHT (4-OHT; Sigma H7904) to the media. Time point 0 was set 12 hrs after the first Hit & Run Cre infection or at the time of 4-OHT addition. Lentiviral particles containing the shRNAs for Ligase 3 (target: CCAGACTTCAAACGTCTCAAA; TRCN0000070978, Sigma), or DNA polymerase theta (target: CGGTCCAACAAGGAAGGATTT; TRCN0000120312, Sigma) in a pLKO,1 vector (Openbiosystem) were produced in 293T cells and introduced into target MEFs with three infections/day (6-12 h intervals) over two days. Infected cells were then transducted with Hit & Run Cre and selected for 3 days in 2-4 μM Puromycin before harvest. Lentiviral particles containing lentiCRISPR v2 with or without sgRNA against NBS1 (GAGAATTACTGTAATCCGCA) were produced in 293T cells and introduced into target MEFs with six infections (4-12 h intervals) over two days after four infections with Hit & Run Cre (6-12 h intervals) over two days. Infected cells were selected for 2 days in 2-4 μM Puromycin before harvest.

**Immunoblotting**

Cells were lysed in 2X Laemmli buffer at 5X10^3^ cells/μl, and the lysate was denatured for 10 min at 95 °C before shearing with an insulin needle. Lysate equivalent to 10^5^ cells was resolved using SDS/PAGE and transferred to a nitrocellulose membrane. Western blot was performed with 5% milk in PBS containing 0.1% (v/v) Tween-20 (PBS-T) using the following antibodies: β-Actin (#3700; Cell Signal); Chk2 (BD 611570; BD Biosciences); DNA-PKcs (SC-1552; Santa Cruz Biotechnology); Ku70 (sc-17789 or sc-1487; Santa Cruz Biotechnology); Lig3 (SC-135883; Santa Cruz Biotechnology); Nbs1 (ab175800; Abcam); TRF2 (#13136; Cell Signal); γ-Tubulin (GTU-88; GeneTex); and secondary anti-Mouse/anti-Rabbit IgG HRP (Cytiva).

Signals were detected according to the manufacturer’s instructions using chemiluminescence western blotting detection reagents (Cytiva) and either BioMax MR film (Kodak) or ChemiDoc (Bio-Rad).

**Chromosome Orientation Fluorescence In Situ Hybridization (CO-FISH) and Immunofluorescence-Fluorescence in situ hybridization (IF-FISH)**

CO-FISH and IF-FISG was performed as previously described ^7^, with minor changes. Briefly, for CO-FISH, cells were labeled with BrdU (7.5μM)/BrdC (2.5μM) for 16 h and treated with 0.2 μg/ml colcemid (Biowest) for one/two hours before collection by trypsinization. Harvested cells were incubated in the hypotonic solution 0.055M KCl at 37 °C for 30 minutes before fixation in MeOH:Ac.Acetic (3:1) overnight at 4 °C. Cells were dropped onto glass slides and allowed to dry overnight. Slides were then rehydrated with PBS, treated with 0.5 mg/ml RNase A (R5000; Sigma) in PBS for 10 min at 37 °C, stained with 1 μg/ml Hoechst 33258 (B2883; Sigma) in 2xSSC for 15 min and exposed to 5.4X10^3^J/m^2^ 365-nm UV light (Stratalinker 1800 UV irradiator). After digestion with 600 U Exonuclease III (M1815, Promega) for 30 min, slides were dehydrated through an ethanol series of 70%, 95%, and 100% and allowed to dry. Staining was performed in hybridization solution (70% formamide, 1 mg/ml blocking reagent (1109617601, Roche), and 10 mM Tris-HCl pH 7.2) with PNA probes from PNA Bio: Cy3-OO-(CCCTAA)_3_ and Alexa Fluor 488-OO-(TTAGGG)_3_, or Alexa Fluor 647-OO-(CCCTAA)_3_ and Cy3-OO-(TTAGGG)_3_. Washes were performed twice in washing solution #1 (70% formamide; 0.1% BSA; 10 mM Tris-HCl, pH 7.2) three times in washing solution #2 (0.08% Tween-20; 0.15 M NaCl; 0.1 M Tris-HCl, pH 7.2) or PBS. To the second wash, DAPI (D1306, Invitrogen) was added to stain DNA. Slides were left to dry and mounted with Prolong Gold Antifade (P36934, Fisher) embedding medium.

For IF-FISH, MEFs were grown on poly-D-Lysine (A3890401, Gibco) pre-coated coverslips for 1-2 days. Cells were rinsed in cold PBS and pre-extracted using cold Triton X-100 buffer (0.1% Triton X-100; 20mM Hepes-KOH, pH 7.9; 50 mM NaCl; 3 mM MgCl_2_; 300 mM sucrose) for 20 minutes on ice, followed by two washes in 1xPBS at RT, before fixation for 10 mins at RT with 3% paraformaldehyde/2% sucrose. Cells were permeabilize for 15 mins with 0.1% Triton X-100 buffer before blocking and staining in Blocking solution (1 mg/ml BSA; 3% goat serum; 0.1% Triton X-100; 1mM EDTA, pH8, in PBS). γH2AX (JBW301, Millipore) primary antibodies, and secondary anti-mouse AlexaFluor 647 antibody (A32728, Invitrogen), were incubated overnight at 4°C or 1 h at RT, respectively. Samples were again fixed in 3% paraformaldehyde/ 2% sucrose for 10 mins at RT before dehydration through the ethanol series of 70%, 95%, and 100% and allowed to dry. Hybridization was performed with Alexa Fluor 488-OO-(TTAGGG)_3_ in hybridization solution (70% formamide; 0.5% blocking reagent (1109617601, Sigma); 10 mM Tris-HCl, pH 7.2) for 10 mins at 45 ^o^C on a heat block, followed by incubation at RT for 2 h, After two washes in washing solution (70% formamide; 10 mM Tris-HCl, pH 7.2), and three in PBS, where DAPI was added to stain the cell nuclei, coverslips were left to dry and mounted with Prolong Gold Antifade embedding medium.

Pictures were acquired on a DeltaVision RT microscope system (Applied Precision) with a PlanApo 60x 1.40 NA objective lens (Olympus America, Inc.) at 1 x 1 binning and multiple 0.2 μm Z-stacks using SoftWoRx software. Images were deconvolved, and 2D-maximum intensity projection images were obtained using SoftWoRx software. Chromatid and chromosome-type fusions were analyzed using Fiji software ^8^ after arbitrary assignment of red for both (TTAGGG) _3_-probes and green for both (CCCTAA) _3_-probes.

Semi-automated analysis and quantification of colocalization was performed using CellProfiler ^9^ with the following pipeline: imaging cropping to remove edge artifacts due to deconvolution; channel intensity rescaling to cover the full histogram range value; “speckle features enhancement” to increase detection sensitivity and remove background/artifact aggregates; “channel-wise primary objects identification” to detect individual nuclei and individual foci; correlation of the foci coordinates in the different channels and with the respective nuclei to define colocalization events. Nuclei with less than 10 detected PNA foci were discarded.

**RNA extraction and qRT–PCR**

RNA was extracted from 10^6^ harvested cells using RNeasy Mini Kit (Qiagen). 500 ng of RNA were reverse transcribed using First Strand cDNA Synthesis Kit (Thermo Scientific). SYBR Green PCR Master Mix (Applied Biosystems) was used for qPCRs. Primers used:

β-actin-F: 5’-TTCTACAATGAGCTGCGTGTGG-3’ ^10^

β-actin -R: 5’-ATGGCTGGGGTGTTGAAGGT-3’ ^10^

PolQ-F: 5’-GCTACCTCCAGAGTCTGTTTCAG-3’

PolQ-R: 5’-ATCCACGACCACCATTCCTAAC-3´

**In-gel analysis of single-stranded telomeric DNA**

Mouse telomeric DNA was analyzed on Clamped homogenous electric field (CHEF) gels as described previously ^7^. Briefly, cells were harvested by trypsinization, resuspended in PBS, mixed with 2% agarose (1:1 ratio) at 50 °C and cast in a plug mold 0.7-1 X 10^6^ cells/plug. Plugs were digested overnight at 50 °C in 1 mg/ml proteinase K (03115879001; Roche) in digest buffer (100 mM EDTA, 0.2% sodium deoxycholate, and 1% sodium lauryl sarcosine) and washed five times in TE. DNA was digested overnight at 37 °C by 60U MboI (#R0147; New England BioLabs). Plugs were then washed in TE, equilibrated in 0.5xTBE and loaded on a 1% agarose/0.5xTBE gel. DNA was resolved by a CHEF-DRII PFGE apparatus (Bio-Rad) for 20 h, with the following settings: initial pulse, 5 s; final pulse, 5 s; 6 V/cm at 14 °C. Gel was dried and hybridized overnight at 50 °C with γ-^32^P-ATP end-labeled TelC (AACCCT)_4_ probe in Church mix (0.5 M sodium phosphate buffer pH 7.2, 1 mM EDTA, 7% SDS, 1% BSA). After three washes in 4xSSC and one in 4xSSC/0.1% SDS at 55 °C, the gel was exposed for one/two days and the single-stranded telomere signal was captured by PhosphoImager. For the acquisition of the total telomere signal, the gel was denatured with 1.5 M NaCl/0.5 M NaOH for one hour, neutralized with two washes of one hour each in 0.5 M Tris-HCl pH 7.0/3 M NaCl, pre-hybridized for 30 min at 55 °C in Church mix, and hybridized overnight at 55 °C with the same probe. The denatured gel was washed and exposed as described before. Quantification of the signals in each lane was done using ImageQuant software. After subtraction of the background, the single-stranded signal was normalized to the total telomeric DNA signal in the same lane. The indicated control value was set to 1, and all the other values were given as a percentage of it.

**Generation and expression of HA-Apollo and Myc-TRF2 mutant alleles**

PCR was used to generate ΔiDDR mutant alleles of Myc tagged mTRF2 (MYC-TRF2) in pLPC retroviral vector using as templates previously published constructs ^11^ and the following primers:

TRF2^ΔiDDR^-Fw: GTTCAGGCACCAGGTGAAGACAG.

TRF2^ΔiDDR^-Rev: TGCTTTGGGCTTCTTCTCCCCCG.

A total number of four infections at 6-12 h intervals were performed before selection for 2-5 days in 2-4 μM Puromycin.

**Protein purification and nuclease assay**

*TRF2*

Human TRF2 and TRF2-ΔiDDR were cloned into a modified pFastbac vector with a His_6_-MBP tag and a 3C protease cleavage site. Proteins were expressed in insect cells grown for 72 hours after infection with baculovirus, harvested, frozen in liquid nitrogen, and stored at -80°C. Cell pellets were thawed and homogenized in lysis buffer (40mM Tris pH 8, 500mM NaCl, 0.5mM TCEP, 10% glycerol, 0.1% Tween-20, 1mM PMSF, protease inhibitor cocktail (Roche)). Cells were pelleted at 18,000x*g* and incubated with 1mL Ni-NTA resin for 1 hr. The resin was washed with A Buffer (40mM Tris pH 8, 100mM NaCl, 0.5mM TCEP, 10% glycerol) and proteins were eluted in A Buffer + 200mM Imidazole. Proteins were incubated with 3C protease overnight at 8°C and injected on a Hi-Trap Heparin column (Cytiva). After washing extensively with A Buffer, proteins were eluted in B Buffer (40mM Tris pH 8, 1M NaCl, 0.5mM TCEP, 10% glycerol). The most concentrated fractions were injected into a Superose 6 column (Cytiva) equilibrated in A Buffer. Fractions were analyzed on SDS-PAGE for purity, pooled, concentrated to 1mg/mL, aliquoted, and flash-frozen in liquid nitrogen.

*MRN/CtIP/DNA-PK*

MRN, CtIP, and DNA-PK (Ku70/80 and DNA-PKcs) were purified as described previously ^12^.

*Endonuclease assay*

The MRN/CtIP endonuclease assay was performed as described ^12^ with the following modifications: MRN was preincubated with the noted concentrations of TRF2 or TRF2ΔiDDR for 10 mins on ice before addition of a separately prepared mixture containing DNA-PK, 5’ ^32^P-labelled substrate DNA, and NU7441. Endonuclease activity was initiated by addition of the remaining components and CtIP and reaction products assessed after 1 hr at 37 °C on polyacrylamide gels containing 12% polyacrylamide, 20% formamide and 6M urea and imaged by phosphorimager.

*Exonuclease assay*

MRN exonuclease assay was performed essentially as described ^13,14^. Briefly, PC1253C (AACGTCATAGACGATTACATTGCTAGGACATCTTTGCCCACGTTGACCCA) was labeled at the 3’ end with alpha-ATP by TdT for 1hr at 37C, heat inactivated at 75°C for 20 minutes, and purified on a G-25 spin column. Labeled oligo (100nM final) was annealed to 2X concentration (200nM final) of PC1253B (TGGGTCAACGTGGGCAAAGATGTCCTAGCAATGTAATCGTCTATGACGTT) with a 3’ Biotin-TEG label. Labeled substrate (1nM final) was pre-incubated with 15nM streptavidin in reaction buffer (16mM Tris pH 8, 40mM NaCl, 4% glycerol, 0.2mM TCEP, 5mM MgCl_2_, 1mM MnCl_2_, 1mM ATP, 0.25mg/mL BSA). Mre11-Rad50 (120nM final) was preincubated with 2.5X Nbs1 (300nM final) and then TRF2 at the indicated concentration was added before adding to the reaction mix. The reaction was incubated at 37C for 2 hours before adding stop buffer (16mM EDTA, 0.3mg/mL Proteinase K, and 0.3% SDS final) and incubating at 50C for 30 minutes. Products were resolved on a 15% TBE-Urea gel (Thermo) and visualized on a Phosphoimager. Three independent replicates were performed and quantified using GraphPad Prism 9.

**AlphaFold-Multimer and Evolutionary and Structural Analysis**

AlphaFold-Multimer (v2.1.0) was run locally on a GPU workstation using default parameters . A custom script (modified from AlphaFold Colab) was used to extract PAE and pLDDT information from the resulting pickle files and structures were analyzed in PyMol (Schrodinger) and ChimeraX (UCSF). TRF2, TRF, CtIP and Rad50 protein sequences were obtained from PSI-BLAST searches of the non-redundant protein sequences (nr) database against human sequences. Alignment of the sequences was performed using MUSCLE in SnapGene and formatted in Jalview. The CtIP alignment was visualized using The NCBI MSA Viewer (v1.22.2) and colored according to “conservation”.

**Quantification and Statistical analysis**

Statistical analysis was performed using GraphPad on three or more independent experiments, as indicated. For each CO-FISH analysis, at least ten metaphases per condition were scored. For in-gel analysis of single-stranded telomeric DNA, the normalized overhang signals were expressed for each cell line independently. Significance was assessed by calculating the p-value using Kruskal-Wallis one-way ANOVA without Gaussian distribution assumption (to compare telomere fusions), unpaired t-test without Gaussian distribution assumption (to compare mRNA expression and telomere overhang signal in a single cell line), and 2-way ANOVA without Gaussian distribution assumption (to compare telomere overhang signal from more than one cell line). P-values ≤ 0.05 were considered statistically significant.

**Acknowledgements**

We are extremely grateful to Devon White for mouse husbandry. Yuming Zhang is thanked for the preliminary experiments with PARPi. Andrea Panza is thanked for help with CellProfiler. Tom Walz and Sarah Cai are thanked for help with AlphaFold-Multimer. This work was supported grants to TdL from the NIH (R35 CA210036 and AG016642) and to FL from Cancerfonden (Can 2018/493) and Vetenskapsrådet (2018-03215). FL is a Wallenberg Molecular Medicine fellow and receives financial support from the Knut and Alice Wallenberg Foundation. BT was partially supported by the Lions forskningsfond (LiU-2022-01245). TP and CV were supported by NIH R01GM138548.

**Conflict of interest**

TdL is a member of the SAB of Calico LLC, San Francisco, CA, USA.

**Extended Data References**

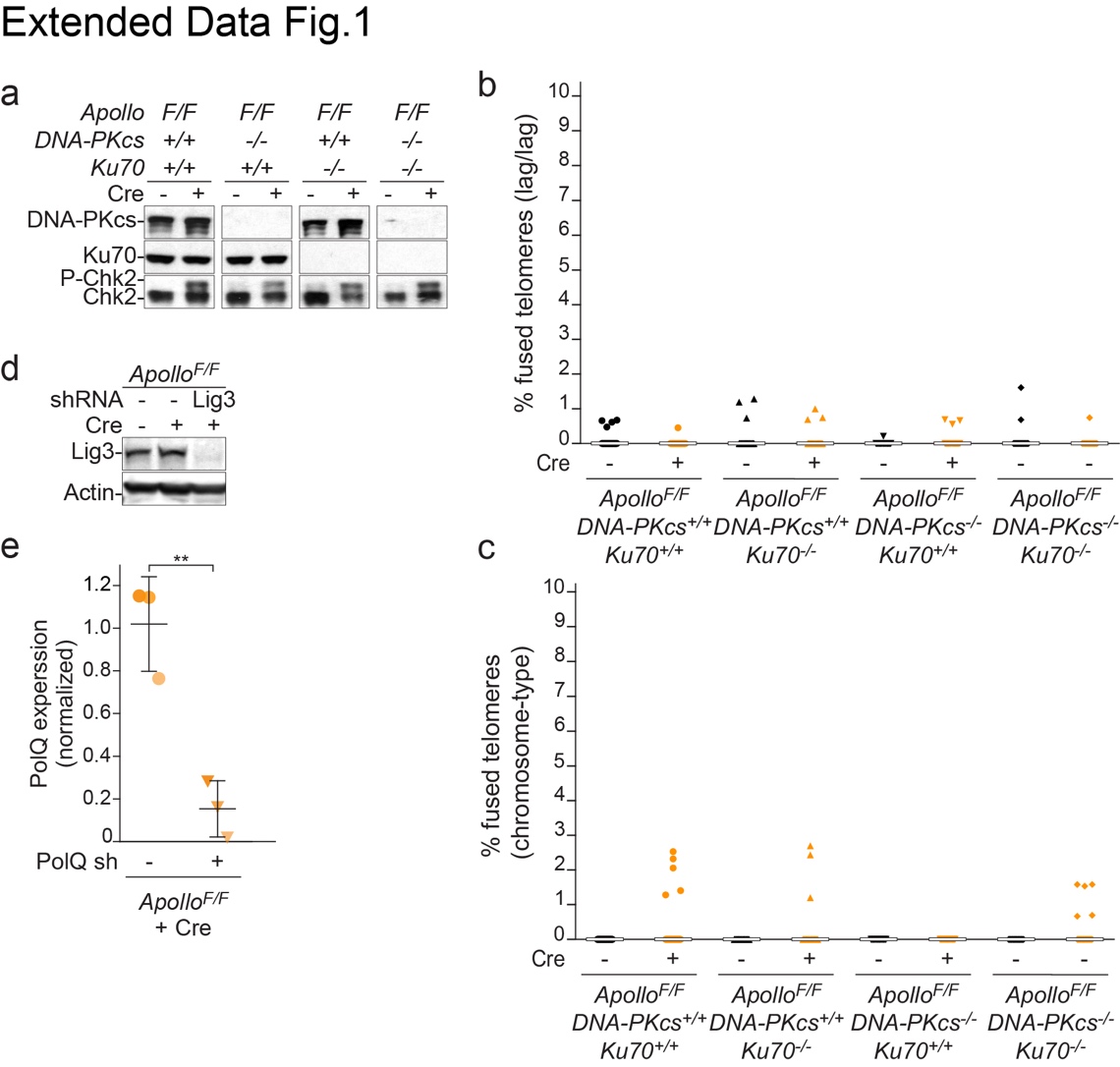

**Extended Data Fig. 1: DNA-PK does not affect Chk2 phosphorylation after Apollo deletion.**

(a) Immunoblots for DNA-PKcs, Ku70, and phosphorylated Chk2 in SV40LT-immortalized Apollo^F/F^, Apollo^F/F^ DNA-PKcs^-/-^, Apollo^F/F^ Ku70 ^-/-^ or Apollo^F/F^ Ku70 ^-/-^ DNA-PKcs^-/-^ MEFs, without any further treatment or 96 h after transduction with Hit & Run Cre, as analyzed in Fig. 1a,b and Fig. 2a,b. (b) and (c) Quantification of telomere fusions as shown in Fig.1a aggregated for chromatid-type involving two lagging-end telomeres (lag/lag) (b) or chromosome-type fusions (chromosome) (c). Bars represent the median. (d) Immunoblots for Lig3 in SV40LT-immortalized Apollo^F/F^ MEFs after transduction with the empty vector or the shRNA against Lig3 and/or 108 h after treatment with Hit & Run Cre as analyzed in Fig. 1g,h. (e) PolQ mRNA expression normalized to β-actin in SV40LT-immortalized Apollo^F/F^ MEFs after transduction with the empty vector or the shRNA against PolQ and 108 h after treatment with Hit & Run Cre, as analyzed in Fig. 1g,h. Values were obtained from three independent experiments and normalized to the empty vectors, with means and SD. Statistical analysis by unpaired t-test with Welch correction.

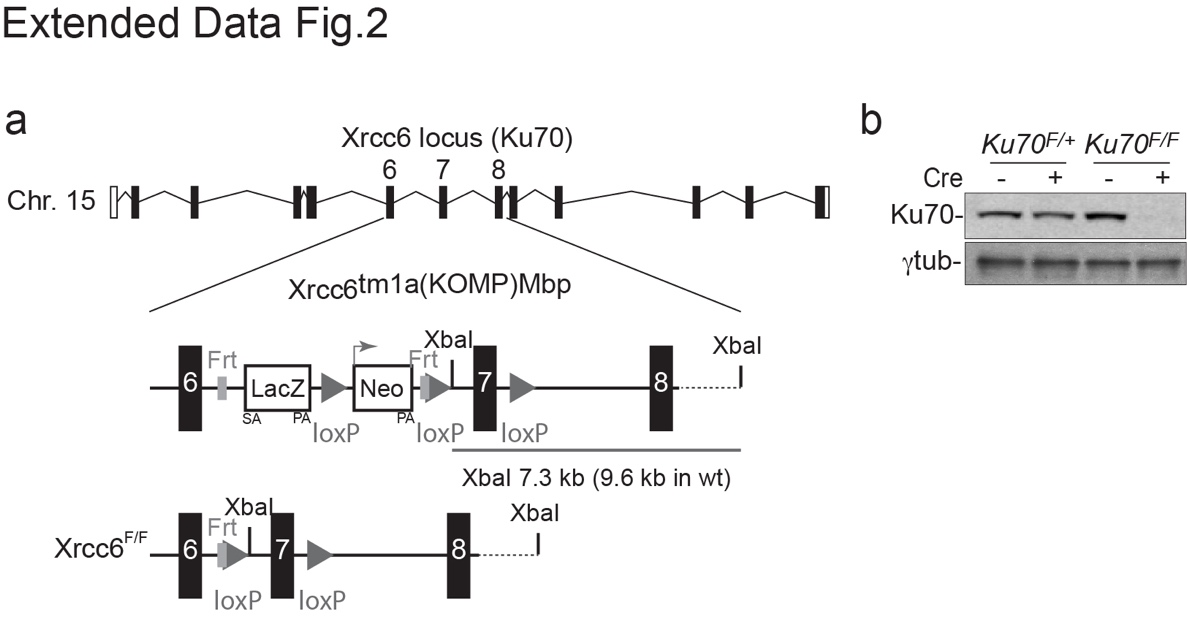

**Extended Data Fig. 2. Conditional deletion of mouse Ku70.**

(a) Targeting of the mouse XRCC6/KU70 locus. The Xrcc6 genomic locus, the KOMP-derived targeted allele with the LacZ/Neo insert and the floxed allele are indicated. The LoxP sites are represented as triangles. (b) Immunoblots for mouse Ku70 in SV40-LT-immortalized *Ku70^F/+^* or *Ku70^F/F^* without any treatment or 108 h after viral transduction with Hit & Run Cre as analyzed in Fig. 2c,d.

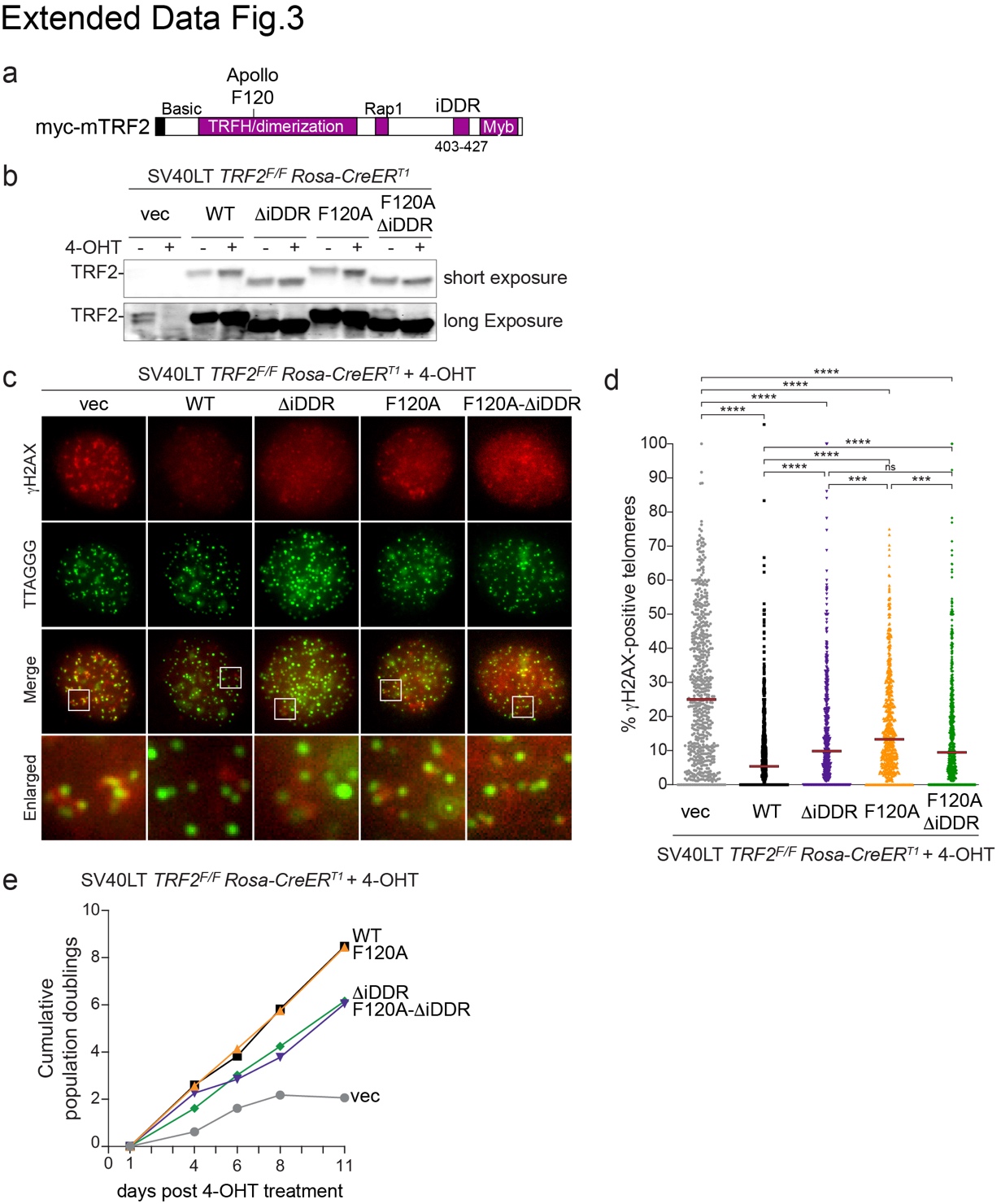

**Extended Data Fig. 3: Expression of MYC-TRF2 alleles.**

(a) Schematic of MYC-tagged mouse TRF2 with Basic, Telomeric Repeat Factors Homology (TRH), Hinge, and Myb domains. Phenylalanine 120 (F120) required for the interaction with Apollo and the iDDR region are highlighted. (b) Lower and higher exposure for the immunoblot of endogenous and exogenous TRF2 as shown in Fig. 4a. (c) IF-FISH of SV40LT-immortalized TRF2^F/F^ RsCre-ERT1 MEFs expressing the empty vector (EV) or the indicated MYC-TRF2 alleles 73 h after 4-OHT-mediated deletion of endogenous TRF2. TIFs are detected by immunofluorescence with antibodies for γ-H2AX (red) and the Telomeres-specific probe Alexa488-OO-(TTAGG)_3_ (green). (d) Quantification of the percentage of TIFs as in (d). Median bars from four independent experiments (150 nuclei per experiment per conditions). Statistical analysis by Kruskal-Wallis Anova test for multiple comparisons. (e) Growth curve showing cumulative population doublings in SV40LT-immortalized TRF2^F/F^ RsCre-ERT1 MEFs expressing the empty vector (EV), in grey, or the indicated MYC-TRF2 alleles: WT in black, ΔiDDR in green, F120A in orange or F120AΔiDDR in purple. 4-OHT was added at time 0. One representative experiment is shown.

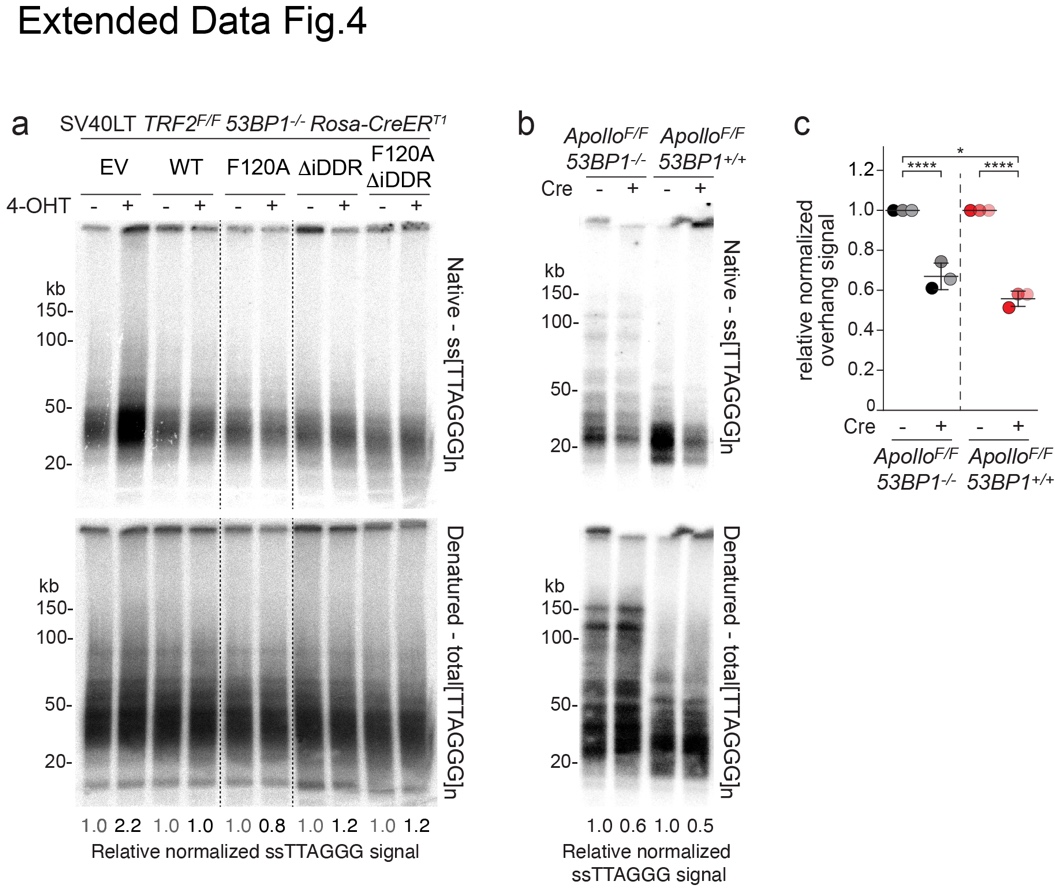

**Extended Data Fig. 4: The iDDR prevents Apollo-independent nucleolytic processing of leading-end telomeres independently from 53BP1.**

(a) Telomeric overhang assay and quantification from one representative experiment in SV40LT-immortalized TRF2^F/F^ 53BP1^-/-^ RsCre-ERT1 MEFs expressing the indicated MYC-TRF2 alleles at 96 h after 4-OH tamoxifen (4-OHT)-mediated deletion of endogenous TRF2. For each allele, the normalized no Cre value was set to 1, and the + Cre value was given relative to it. (b) and (c) Telomeric overhang assay and quantification of Apollo^F/F^ and Apollo^F/F^ 53BP1^-/-^MEFs 96 h after Hit & Run Cre-mediated deletion of Apollo as described in (a) and (b), with means and SDs across three independent experiments. Statistical analysis by two-way ANOVA.

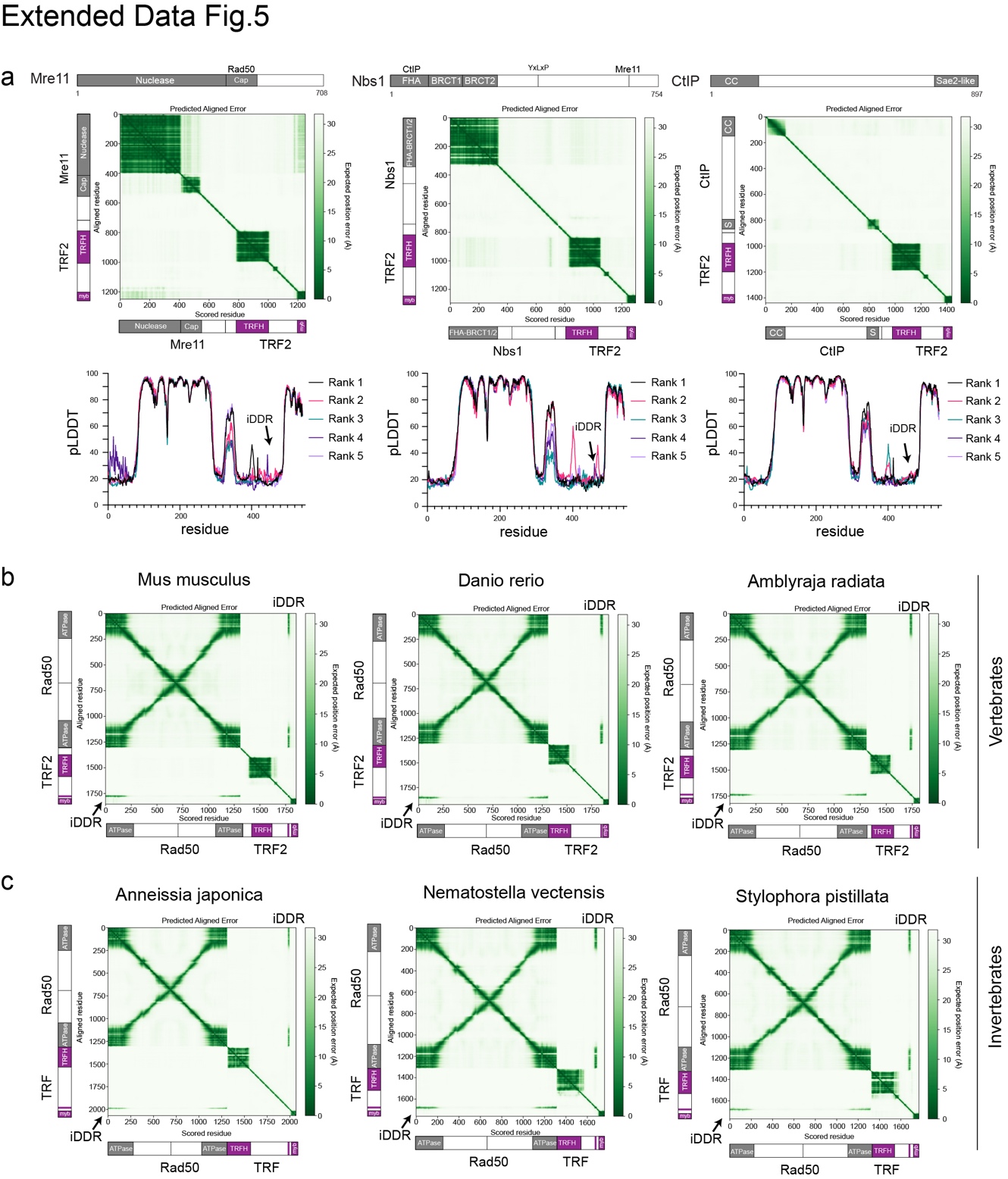

**Exteded Data Fig. 5: The iDDR is predicted to interact with Rad50 in several metazoan.**

(a) Representative Predicted Aligned Error (PAE; top) of TRF2 with Mre11 (left), Nbs1 (middle), and CtIP (right) from AlphaFold-Multimer models and predicted local distance difference test (pLDDT; bottom) for each residue in TRF2 from five ranked models generated with default parameters. (b) Representative Predicted Aligned Error (PAE) for the interaction between TRF2 and Rad50 in representative vertebrates. (c) Representative Predicted Aligned Error (PAE) for the interaction between TRF and Rad50 in representative invertebrates.

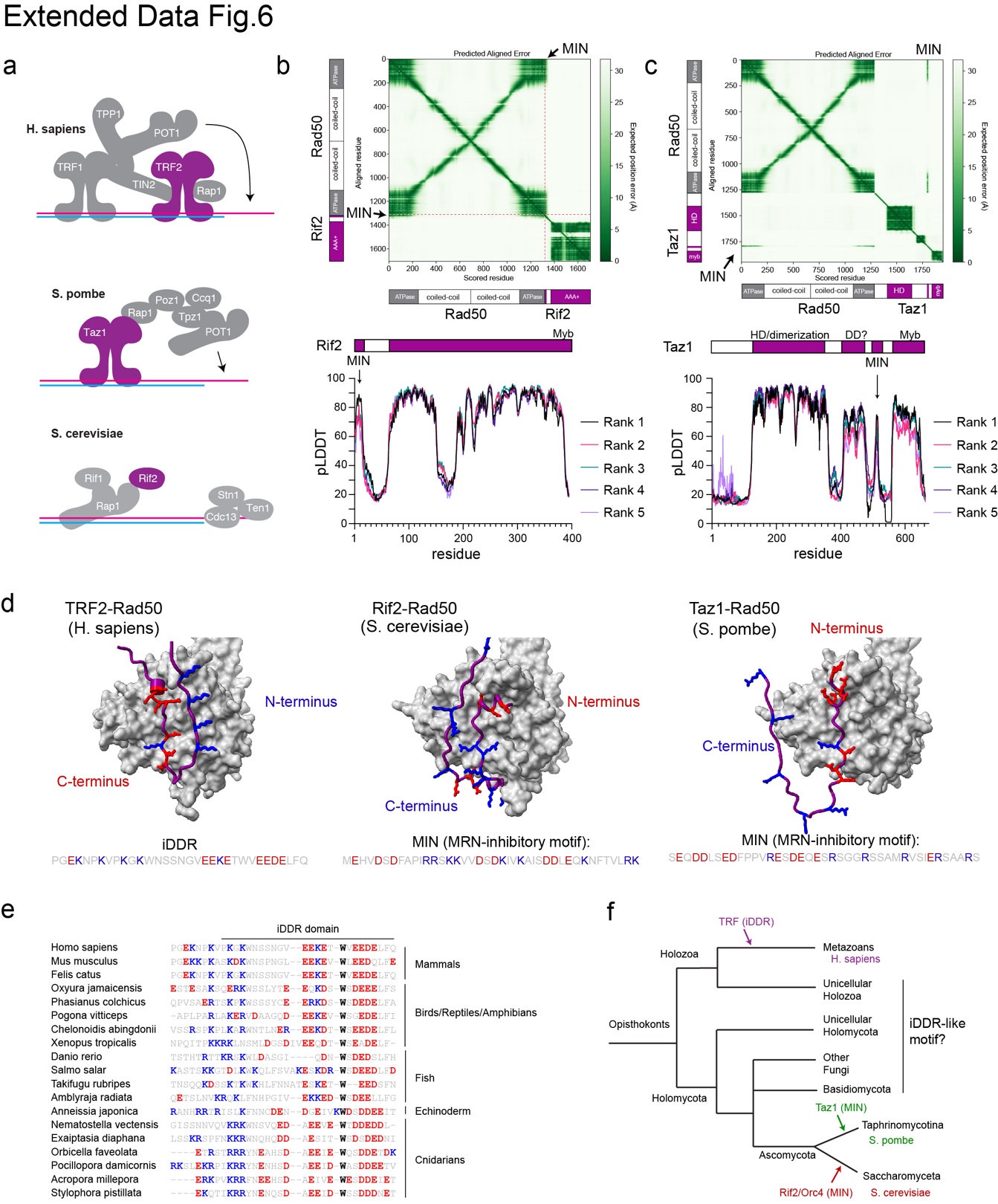

**Extended Data Fig. 6: The iDDR of TRF2 and the MIN domains of Rif2 and Taz1 are an example of convergent evolution.**

(a) Schematic for telomere binding proteins in *H. sapiens*, *S. pombe*, and *S. cerevisiae* with TRF2, Taz1, Rif2 highlighted in purple. (b) AlphaFold-Multimer Predicted Aligned Error (PAE; top; representative of five ranked models generated with default parameters) plot for S. cerevisiae Rif2 and Rad50 and predicted Local Distance Difference Test (pLDDT; bottom) plot for each residue in Rif2 from the ranked models.(c) AlphaFold-Multimer PAE (top; representative of five ranked models generated with default parameters) plot for S. pombe Taz1 and Rad50 and pLDDT (bottom) plot for across Taz1 from the ranked models. (d) Representative AlphaFold-Multimer models of TRF2-Rad50 (left), Rif2-Rad50 (middle), and Taz1-Rad50 (right). Acidic (red) and basic (blue) residues are highlighted. (e) MUSCLE alignment of the iDDR domains from representative metazoans, highlighting the presence of basic residues (blue) followed by acidic residues (red). (f) Phylogenetic tree of Opisthokonts showing the emergence of the iDDR of TRF proteins (blue) as well as the MIN of Taz1 (green) and the MIN (or BAT) of Rif2/Orc4 (red). Whether other Opisthokonts independently evolved an iDDR-like motif is unknown. Not to scale.

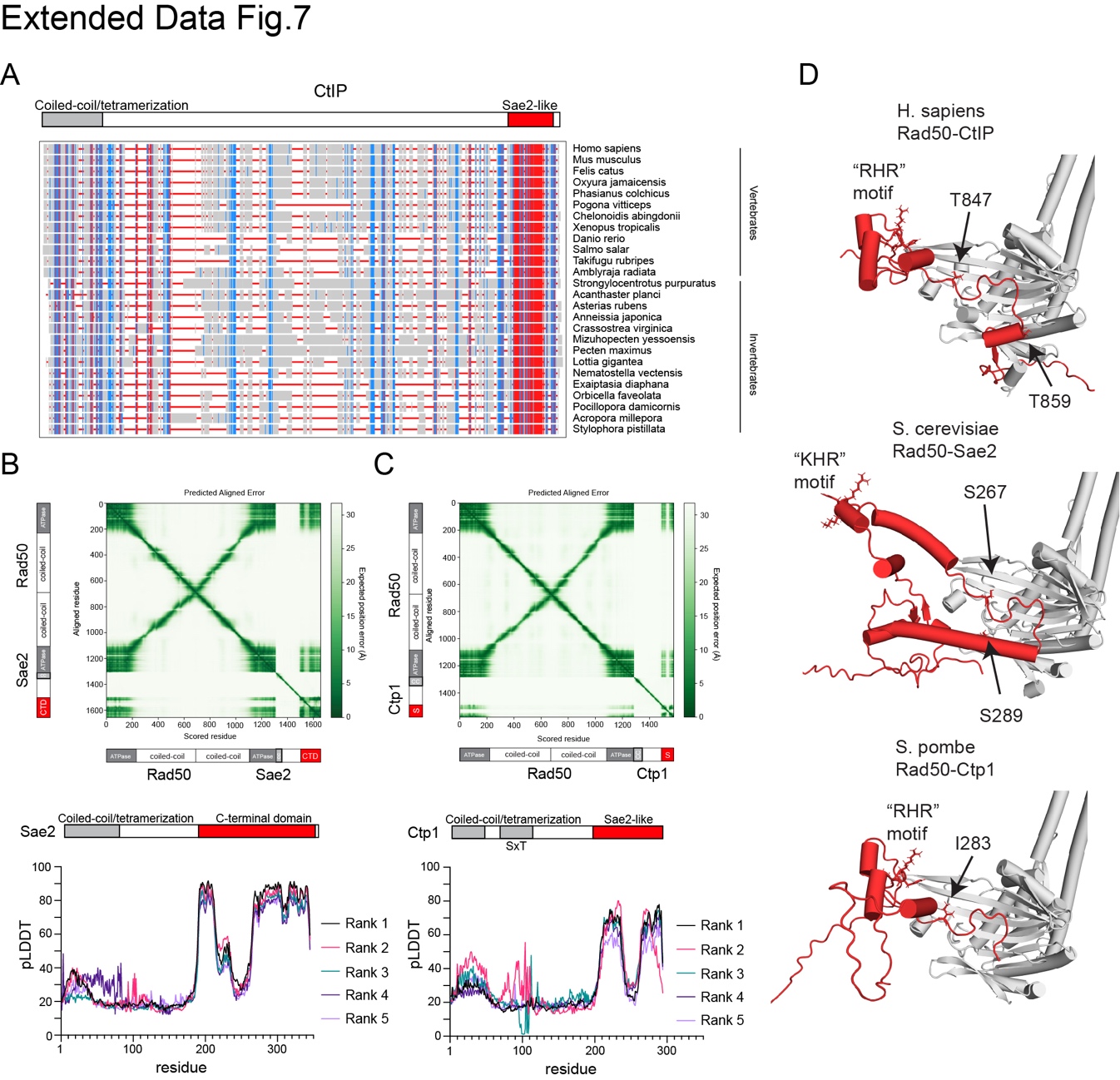

**Extended Data Fig. 7: CtIP, Sae2, and Ctp1 are all predicted to interact with Rad50 in a similar manner.**

(a) Multiple Sequence Alignment (MSA) of CtIP proteins from vertebrate and invertebrate species using NCBI MSA Viewer from an alignment using Multiple Sequence Comparison by Log-Expectation (MUSCLE). Vertical lines are colored by conservation where red indicates highly conserved and blue indicates lower conservation. Alignment positions with gaps are not colored. (b) Representative AlphaFold-Multimer models for CtIP-Rad50, Sae2-Rad50, and Ctp1-Rad50. Important CDK and ATM/Tel1 sites are indicated as well as the residue in the ATM site position on Ctp1. (c) AlphaFold-Multimer Predicted Aligned Error (PAE; top; representative of five ranked models generated with default parameters) plot for S. cerevisiae Sae2-Rad50 and predicted Local Distance Difference Test (pLDDT; bottom) plot across Sae2 from the ranked models. (d) Representative AlphaFold-Multimer models for CtIP-Rad50, Sae2-Rad50, and Ctp1-Rad50. Important CDK and ATM/Tel1 sites are indicated as well as the residue in the ATM site position on Ctp1.
